# Reward changes the use and structure of predictive spatial maps

**DOI:** 10.64898/2026.10.28.762439

**Authors:** Sharvil Oza, Amir Cohen, Genevieve Wager, Dori Derdikman, Stefano Recanatesi

**Affiliations:** Dhirubhai Ambani University, Gandhinagar, India; Technion — Israel Institute of Technology, Haifa, Israel; Allen Institute for Neural Dynamics, Seattle, WA, USA

## Abstract

Reward-related changes in hippocampal place-cell activity may reflect either different use of an existing spatial map or plasticity that reshapes the map. We separated these contributions in a two-pathway model: a recurrent network learned to predict sensory input and developed a spatial representation, while a reinforcement-learning controller combined this representation with an explicit goal signal to guide behavior. Replaying identical trajectories before and after learning isolated representational plasticity. Goal-directed behavior strongly concentrated place fields near reward, whereas changes to the representation were smaller and more variable. A reward-proximity teaching signal made these changes larger and more consistent, and the change was strongly attenuated when recurrent plasticity was restricted. After reward relocation, goal-related directional tuning adapted more fully than place-field organization, while withdrawing reward and proximity teaching reversed the learned field concentration. Thus, reward affects predictive spatial maps through separable mechanisms: goal information changes how an existing map is used, whereas reward-related teaching signals reshape the map itself. Our results provide a concrete framework for future experiments to disentangle how reward information, plasticity and behavior jointly shape hippocampal activity.

---

Hippocampal place cells, neurons active at specific locations in an environment (1), form a spatial representation shaped by experience, environmental features, context and behavioral demands (2–4). Reward-related changes in this representation can arise through at least two routes. A change in policy alters the trajectories, headings and sensory histories through which an existing map is sampled and expressed, whereas reward-dependent learning can modify the map itself through plasticity. Place fields can accumulate around rewarded locations (5–11) and shift toward learned goals (12, 13), while goal-related firing can also occur outside a cell’s primary field (14). These effects take multiple forms and are task-dependent (15, 16): goal over-representation was absent when goal location was dissociated from reward delivery and in a rapid-learning paradigm in which goal information emerged instead during ripples (17, 18). However, changes measured during goal-directed behavior do not by themselves distinguish a different *use* of an existing map from reward-driven *reshaping* of the representation.

Predictive accounts offer one framework for hippocampal spatial coding. The successor representation describes states by their expected future occupancy under a policy (19– 22), while recurrent networks trained to predict sensory input can develop place-cell-like tuning without an explicit spatial objective (23–26). Hippocampal reward representations can also become predictive with learning, including activity that progressively shifts from reward delivery toward earlier reward-predicting states (27). These findings link prediction, spatial coding and reward, but do not establish how reward-dependent plasticity acts on a predictive spatial representation.

This distinction is particularly important for recurrent predictive maps. Even without changing the underlying parameters, a new policy changes the sequences of states and sensory inputs encountered during behavior and can therefore change the spatial activity that is observed. Plasticity can independently alter the recurrent representation, causing the same trajectory to evoke different activity after learning. Observed remapping can thus combine changes in how a map is *used* with changes in how it is *rewritten*. A model that separates these contributions is needed to determine when reward changes the expression of a predictive map and when it changes the map itself.

The successor representation makes this ambiguity explicit: because future occupancy depends on policy, changing behavior changes the representation it predicts even when the environment is unchanged, including an apparent clustering of fields near frequently visited rewards (19–21). Thus, differences between rate maps measured under different policies can reflect behavioral sampling, learning-dependent changes in the underlying representation, or both.

To separate these contributions, we constructed a two-pathway model in the spirit of earlier accounts that pair a hippocampal spatial representation with a reinforcement-learning controller (28). A recurrent network learned to predict the next egocentric sensory observation and developed a spatial representation, while a downstream actor–critic combined this representation with a supplied allocentric displacement to the goal to select actions. We then replayed the same sensory–action trajectory through the recurrent network before and after reward learning. Holding position, heading and sensory history fixed isolates changes in recurrent activity caused by learned parameters from those associated with a change in policy. Goal-directed behavior produced a strong concentration of spatial activity near reward, whereas fixed-trajectory replay revealed a smaller and more variable representational change. Adding an auxiliary reward-proximity objective made this change larger and consistently directed toward reward. Thus, goal-directed readout and reward-dependent plasticity make separable contributions to the spatial activity observed during behavior.

The model is a computational abstraction rather than a circuit model of hippocampal reward learning. It assumes an explicit goal-displacement signal rather than learning one from experience. Goal representations in orbitofrontal cortex and hippocampal sequences and vector fields directed toward remembered or current goals support the availability of goal information during navigation (16, 29–33), but do not specify the source or coordinate system assumed here. The model also receives an externally defined reward-proximity teaching signal. Although spatially related signals have been observed in dopaminergic inputs to hippocampus (34), this target should not be interpreted as dopamine activity or a reward-prediction error. Nor does the model reproduce the trial-by-trial backward progression of reward-related activity reported during learning (27).

Within these limits, the model reproduces several features of hippocampal reward coding without being fitted to a single dataset: the concentration and movement of place fields toward reward (5, 6, 11–13), the loss of reward-centered organization following reward omission (34), and the coexistence of place-anchored and reward-relative responses after reward relocation (35). We further compare the model with hippocampal recordings from reward-omission and reward-relocation experiments (34, 35). Together, these analyses provide a framework for distinguishing changes in how a predictive spatial map is used from changes in the representation produced by reward-dependent learning.

## Results

### Sensory prediction establishes a spatial representation that can be recruited for goal-directed navigation

Can a spatial representation acquired during exploration subsequently support navigation to reward? We first considered how such a representation could emerge without a behavioral goal. An agent explored a square arena while receiving egocentric visual features capturing what lay ahead, and a recurrent network learned to predict the next sensory observation from the current observation and action (Fig. 1; Fig. 2a) (26). During this initial learning period, termed Phase A, the network received no reward, target or explicit position signal. Sensory prediction thus provided the sole learning objective.

**Fig. 1.**
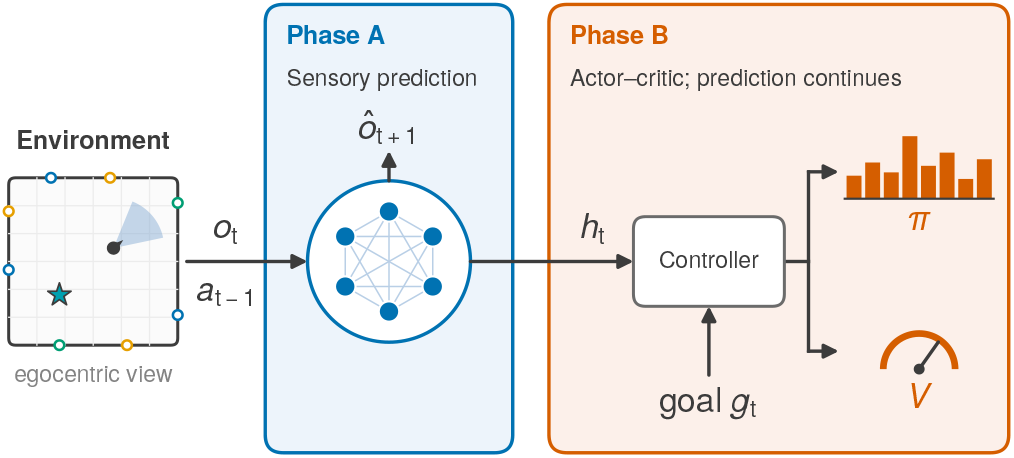
Predictive recurrent network and goal-conditioned controller. In Phase A, a recurrent network learns to predict the next egocentric observation from the current observation and outgoing action. During online control the recurrent update instead receives the preceding action, as detailed in Methods. In Phase B, an actor–critic combines the recurrent state with an allocentric goal displacement to select actions, while sensory prediction continues.

**Fig. 2.**
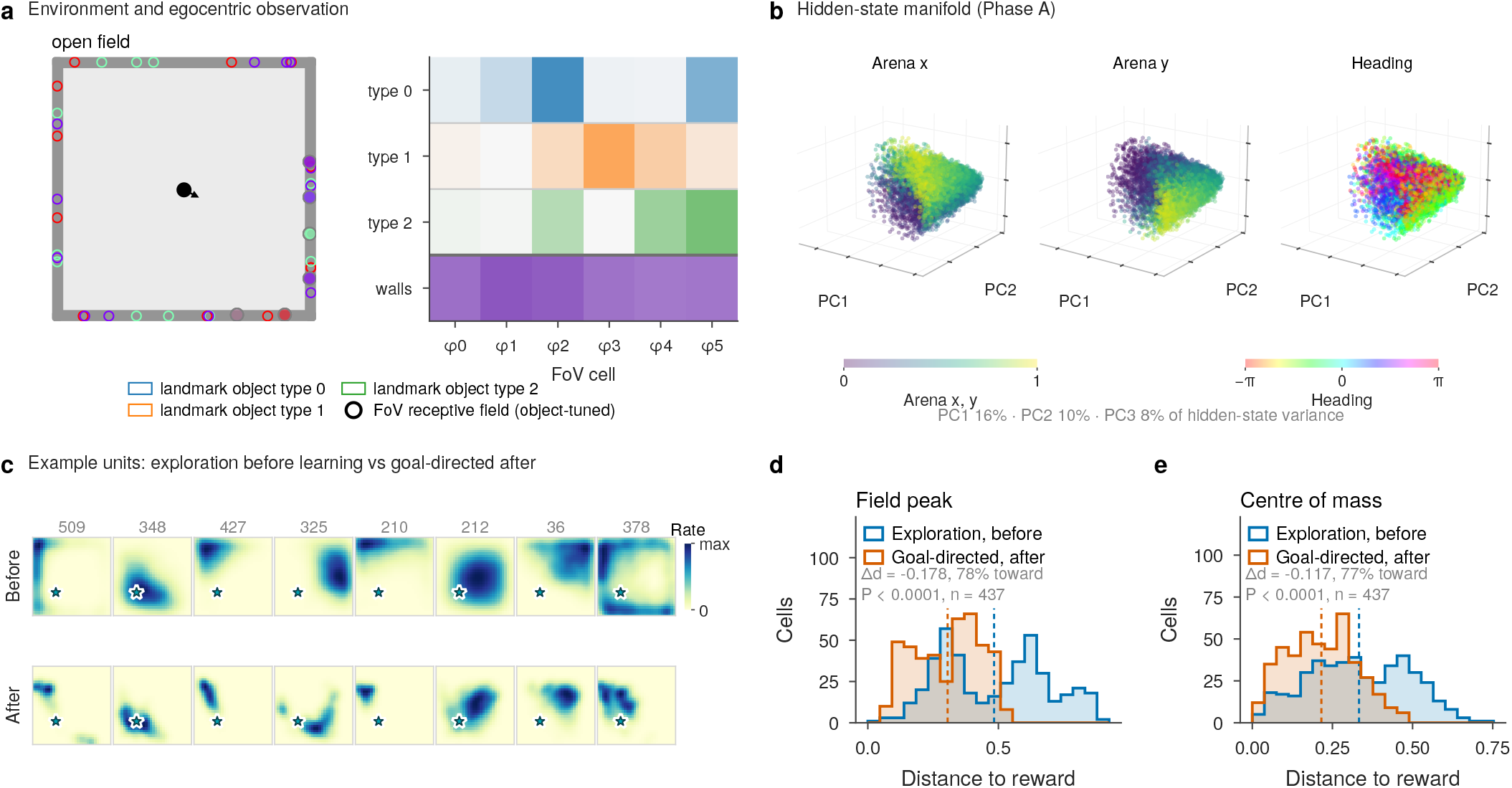
Sensory prediction produces a spatial representation that supports goal-conditioned control. Phase A sensory prediction yields place-cell-like spatial tuning, whose fields are compared before and after Phase B reward learning. a) The environment and one observation. Each timestep supplies 24 egocentric sensory values: 18 field-of-view cells (six for each of three landmark object types), plus six wall-distance cells, and no allocentric position. b) Three-dimensional principal-component projection of Phase A (exploration before any reward) population activity, colored by arena position and by heading. c) Example units of one network (close to the cohort median) during exploration with no reward present (top) and along goal-directed trajectories after reward learning (bottom); the star marks the reward location. d,e) Distance to reward (arena units; arena is 1 *×* 1) of the field peaks (d) and centers of mass (e) of the same example network’s place cells (*n* = 437 units with Skaggs information ≥ 0.05 bits/spike in both maps), before learning (exploration) and after learning (goal-directed behavior). Both the network parameters and the sampled trajectories differ between the two, so this contrast mixes learning with behavior. Dashed lines, means; annotations give the mean per-cell change (negative = toward reward), the percentage of cells moving toward reward and a paired Wilcoxon test. Statistics across all 55 retained networks are reported in the text and Fig. S1.

Through this learning, recurrent population activity acquired a spatial organization. Hidden states occupied a low-dimensional manifold whose geometry reflected the arena, with nearby physical positions represented by nearby population states (Fig. 2b). Individual units developed compact spatial fields that varied in size, shape and location and approximately tiled the environment (Fig. 2c, top row). These place-cell-like responses emerged before reward was introduced, without a systematic bias in field centers. Sensory prediction therefore established a representation of space during exploration.

We next asked how this existing representation would be used and modified when behavior became directed toward reward. In Phase B, we added a downstream action-selection pathway, implemented as an actor–critic controller, and trained it to guide the agent to rewarded locations (Fig. 1). Action selection combined a readout of the recurrent population state with an explicitly supplied allocentric displacement to the current goal. The goal signal was therefore available to the control pathway rather than inferred by the predictive network. The recurrent network continued to learn through sensory prediction, allowing its parameters to adapt to the sensory sequences encountered during goal-directed behavior.

As the controller learned to reach reward, the recurrent representation retained its spatial organization. Measured on an identical replayed trajectory, most units remained place cells at the end of Phase B (71.5% of units, s.e.m. 1.8, versus 80.7%, s.e.m. 1.0, at the end of Phase A), and their spatial information increased (2.75 versus 1.73 bits, s.e.m. 0.15 and 0.03; Fig. S1a,b). Nevertheless, spatial fields measured along the resulting goal-directed trajectories were more concentrated near reward (Fig. 2c–e). Comparing exploration before learning with goal-directed behavior after learning, across the 55 networks retained for this comparison (see Methods), the distance of field peaks to reward decreased by 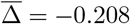 (s.d. 0.040, s.e.m. 0.005; *p* = 1.1 × 10^−10^), with an average of 83.8% of cells per network shifting toward reward and a decrease in every network. Field centers of mass gave the same result (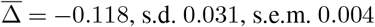; 76.6% of cells; *p* = 1.1×10^−10^; Fig. S1c–f).

Thus, a representation acquired through sensory prediction could be recruited for goal-directed navigation, during which its measured spatial fields became biased toward reward. Because both the learned parameters and the trajectories used to sample activity differed between phases, this comparison alone could not establish how much of the bias reflected a lasting change in the representation. We therefore next held the complete evaluated trajectory fixed and asked how much of the reward-related concentration remained when only the learned parameters differed.

### Fixed-trajectory replay reveals a parameter-dependent shift toward reward

How much of the concentration of spatial fields near reward reflected changes in the learned representation? Comparing rate maps across exploratory and goal-directed behavior cannot resolve this question, because both the network parameters and the trajectories used to measure activity differ. For a recurrent network, matching spatial occupancy alone is insufficient: activity at a given location also depends on the preceding sensory and action sequence. We therefore compared the pre- and post-learning networks using an identical observation–action trajectory (Fig. 3).

**Fig. 3.**
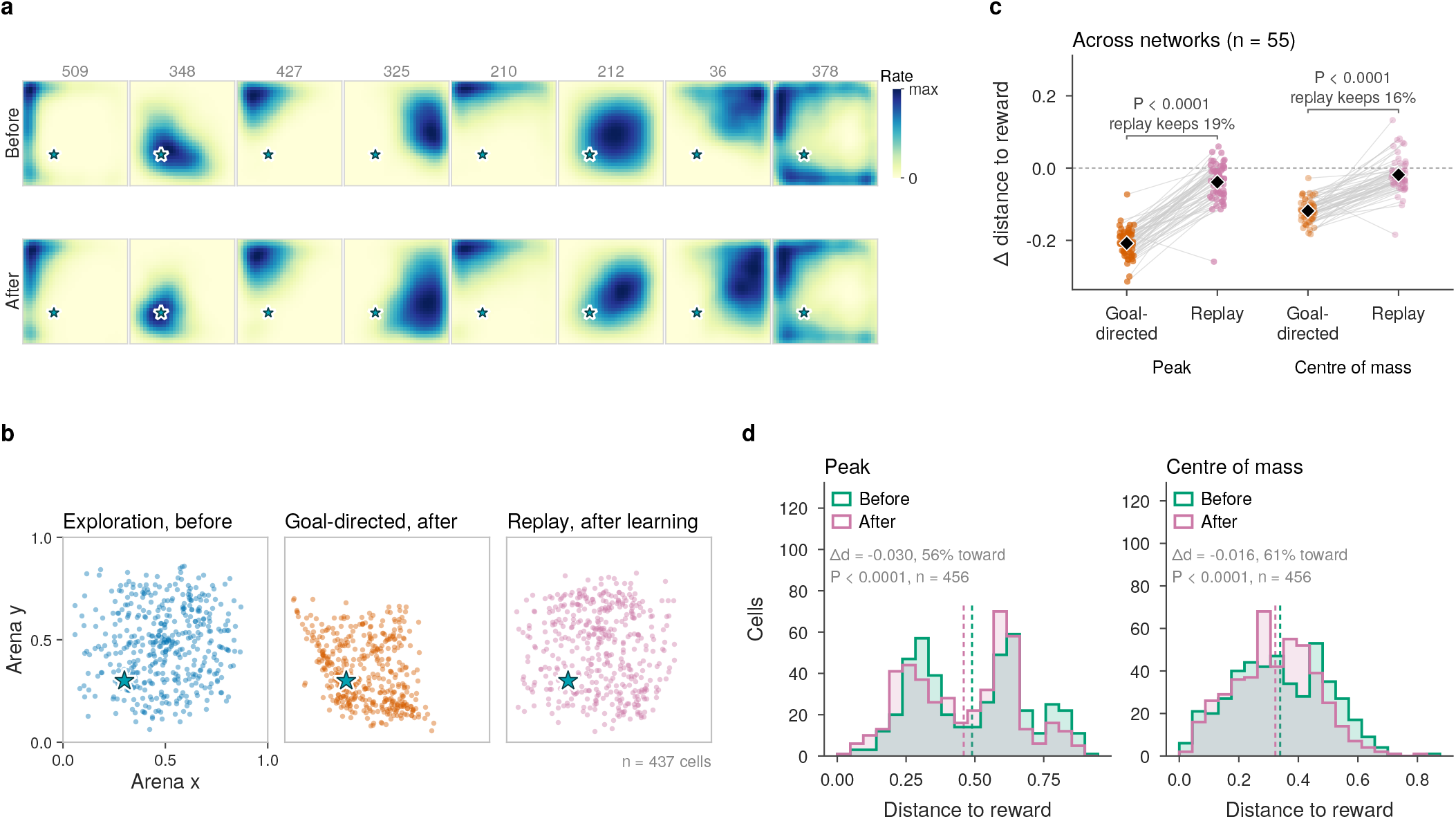
Fixed-trajectory replay separates goal-conditioned readout from parameter-dependent reorganization. Fixed-trajectory replay passes the same stored exploratory observation–action trajectory through the network before and after reward learning, so differences reflect learned parameters alone. Most of the shift toward reward seen during goal-directed behavior disappears under replay. a) Fields of the example network’s units (same units as Fig. 2c; star = reward) before and after learning. b) Field centers of mass of the example network’s place cells (each dot is one cell; *n* = 437, Skaggs information ≥ 0.05 bits/spike in both the exploration and goal-directed maps) during exploration before learning, along goal-directed trajectories after learning and on the replayed exploratory trajectory after learning. c) Per-network mean change in distance to reward (arena units; negative = toward reward) from exploration before learning to goal-directed behavior after learning (parameters and trajectory differ) and under fixed-trajectory replay (only parameters differ), for field peaks and centers of mass, in the 55 retained networks. Gray lines join each network’s two estimates; diamonds, mean *±* s.e.m.; brackets, paired Wilcoxon tests and the fraction of the behavioral shift retained under replay: 19% for peaks, 16% for centers of mass. d) Distance to reward of the example network’s place cells on the replayed trajectory before and after learning, for peaks and centers of mass (*n* = 456 cells with Skaggs information ≥ 0.05 bits/spike in both the before and replay maps; dashed lines, means; annotations give the mean per-cell change, the percentage of cells moving toward reward and a paired Wilcoxon test).

Before Phase B, we recorded a long exploratory trajectory and used the associated observations and actions to construct baseline fields. After reward-driven learning (Phase B), we replayed this sequence through the updated network, holding position, heading, actions and sensory history fixed. This *fixed-trajectory replay* isolated differences associated with the learned parameters from those arising from changes in the evaluated trajectory. Paired rate maps from the example network of Fig. 2 showed that individual fields remained largely similar, with modest shifts toward reward (Fig. 3a). Like-wise, field centers measured along the replayed exploratory trajectory remained closer to the exploratory distribution than those measured during goal-directed behavior (Fig. 3b).

Across the 55 networks, most of the shift measured during goal-directed behavior disappeared under fixed-trajectory replay (Fig. 3c). Field peaks moved by 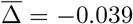 (s.d. 0.053, s.e.m. 0.007) under replay, compared with −0.208 between exploration before learning and goal-directed behavior after it; replay retained 19% of the behavioral shift (median per network 16%), and the replay shift was smaller in 54 of 55 networks (paired *p* = 1.3 × 10^−10^). Centers of mass behaved alike: −0.019 (s.d. 0.050, s.e.m. −0.007) under replay against 0.118 during behavior, 16% retained (paired *p* = 1.0 × 10^−9^). The residual replay shift was nevertheless consistent: 57.1% of cells moved toward reward and the network mean was negative in 42 of 55 networks (*p* = 2.7 × 10^−6^; centers of mass, 58.1% of cells, *p* = 1.9 × 10^−3^), and in the example network the distance distributions shifted modestly toward reward (Fig. 3d).

Thus, the concentration of fields near reward measured during goal-directed behavior exceeded, by about fivefold, the change expressed by the learned network along a fixed trajectory. Learning produced a modest parameter-dependent bias toward reward, while most of the behavioral contrast depended on the trajectories along which activity was measured. We next asked what determines the magnitude and direction of this parameter-dependent reorganization.

### A reward-proximity teaching signal directs plastic reorganization toward reward

What determines if changes in the spatial representation are directed toward reward? We examined two factors: plasticity (i.e., the capacity of recurrent connections to change) and the reward-related information available to guide that change. To manipulate reward teaching content, we added an auxiliary readout trained to predict a smooth function of Euclidean distance to the current reward (Methods). Errors in this prediction updated the recurrent parameters, encouraging the population state to encode reward proximity. The target was externally specified and served as a phenomenological teaching signal; it did not model how dopamine activity is generated. It also differed from the controller’s critic, which estimated policy-dependent discounted return.

With reward-proximity teaching, fixed-trajectory replay revealed a larger and more consistent shift toward reward. In a network whose population shifts were close to the cohort means, selected well-tuned fields illustrate this movement, and the full population of field centers moved toward the rewarded location despite identical evaluated trajectories (Fig. 4a,b). Across the 55 networks meeting the inclusion criteria (Fig. 4c), field peaks moved toward reward by 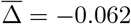 (s.d. 0.032, s.e.m. 0.004; 65.3% of cells; *p* = 2.4 × 10^−10^) and centers of mass by 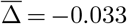 (s.d. 0.027, s.e.m. 0.004; 63.1% of cells; *p* = 1.1 × 10^−8^). The mean field-peak shift was toward reward in 54 of 55 networks, and its across-network standard deviation relative to the magnitude of the mean decreased from 1.37 without proximity teaching to 0.51 with it. The example network’s cell-level distributions shifted accordingly on both measures (Fig. S2c). The result persisted across entropy-gate and place-cell information thresholds (Fig. S2a,b).

**Fig. 4.**
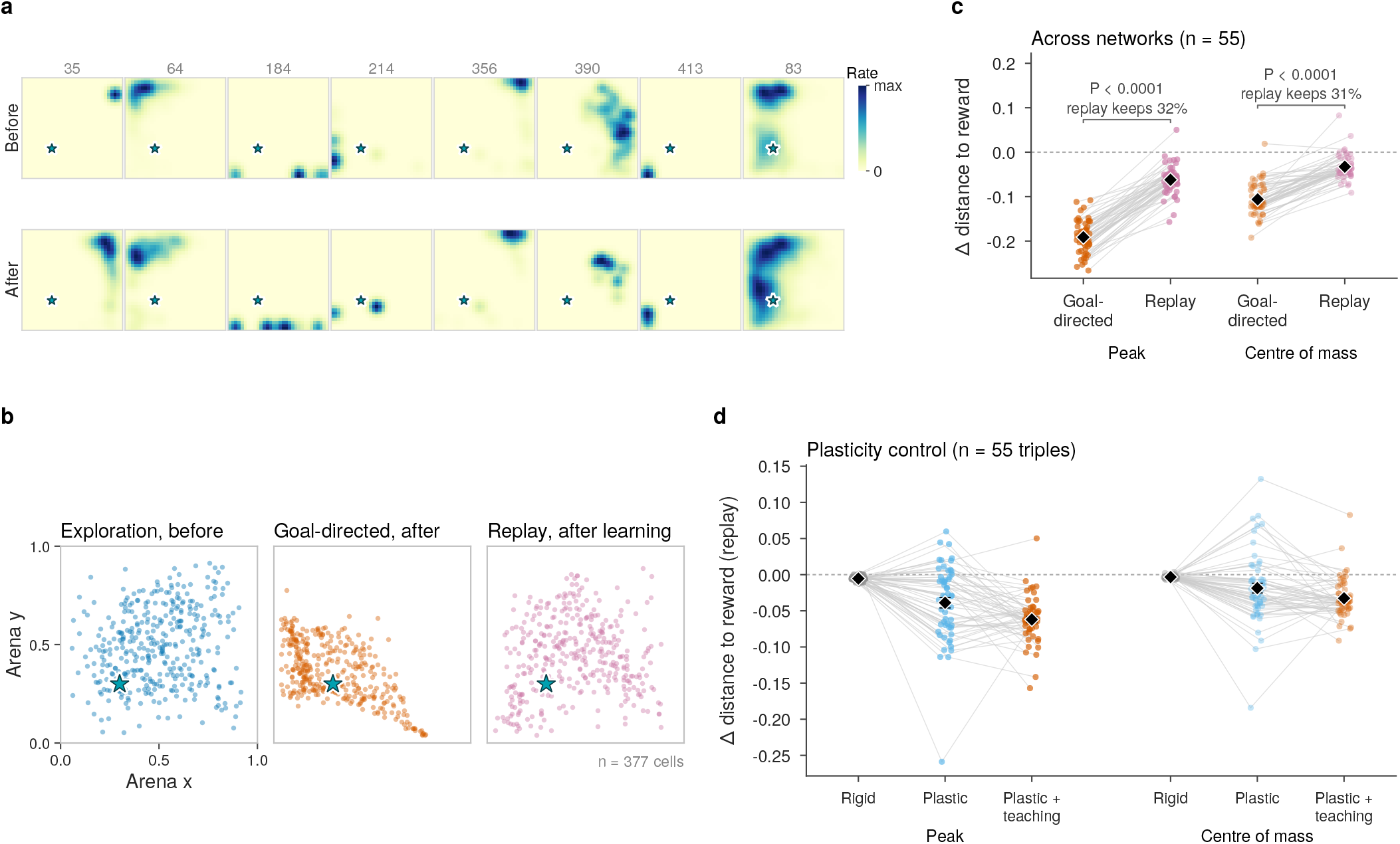
A reward-proximity teaching objective directs plastic reorganization toward reward. During reward learning, an auxiliary readout of the recurrent state is additionally trained to predict a reward proximity target: a smooth, externally specified value that is highest at the reward and falls with distance from it. The prediction error also updates the recurrent weights (Methods). Fixed-trajectory replay passes the same stored exploratory trajectory through the network before and after learning. a) Fields of selected units of the example network (star = reward) before and after learning. b) Field centers of mass of the example network’s place cells (*n* = 377, Skaggs information ≥ 0.05 bits/spike in both maps; each dot is one cell) during exploration before learning, along goal-directed trajectories after learning and on the replayed trajectory after learning. c) Per-network mean change in distance to reward (arena units; negative = toward reward) from exploration before learning to goal-directed behavior after learning, and under fixed-trajectory replay, for field peaks and centers of mass, in the 55 retained networks with proximity teaching (gray lines join a network’s two estimates; diamonds, mean *±* s.e.m.; brackets, paired Wilcoxon tests and the fraction of the behavioral shift retained under replay: 32% for peaks, 31% for centers of mass). d) Matched plasticity control for field peaks and centers of mass under fixed-trajectory replay. Each gray line joins a triple of networks with identical pre-learning states: near-rigid recurrent weights, plastic recurrent weights without proximity teaching and plastic recurrent weights with proximity teaching (*n* = 55 triples); diamonds show mean *±* s.e.m.

Matched network triples directly separated the roles of recurrent plasticity and teaching content (Fig. 4d). The three networks in each triple began from an identical pre-learning state, differed only in recurrent plasticity or proximity teaching during reward learning, and were evaluated on the same replay trajectory. Restricting recurrent plasticity reduced the mean field-peak shift toward reward by approximately an order of magnitude, to 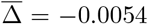 (s.d. 0.0034, s.e.m. 0.0005; *p* = 4.3 × 10^−10^). Mean absolute peak displacement was also reduced, to 0.028 (s.d. 0.013, s.e.m. 0.002), compared with 0.315 (s.d. 0.076, s.e.m. 0.010) in the plastic condition without proximity teaching and 0.286 (s.d. 0.072, s.e.m. 0.010) with it. Among the plastic networks, proximity teaching increased the mean field-peak shift toward reward from −0.039 to −0.062 (*p* = 1.8 × 10^−3^), with a larger shift in 67% of matched pairs. The center-of-mass comparison supported the same conclusion (−0.019 to −0.033; *p* = 0.013). Because peaks are defined on spatial bins and can switch between sub-fields, absolute peak displacement measures changes in peak location rather than continuous movement of individual fields. These comparisons distinguish the extent of reorganization from its direction: recurrent plasticity permitted substantial changes in field location, whereas proximity teaching increased their component toward reward without increasing total peak displacement.

Finally, we asked whether field concentration during goal-directed behavior captured this effect of teaching content. In the 55 matched pairs, the shift measured between exploration before learning and goal-directed behavior after learning was larger without proximity teaching than with it (−0.208 versus −0.191; *p* = 6.6 × 10^−3^). Fixed-trajectory replay gave the opposite ordering, revealing a larger shift with proximity teaching (−0.039 versus −0.062; paired *p* = 1.8 × 10^−3^). With proximity teaching, replay retained a larger fraction of the behavioral shift (32% for peaks and 31% for centers of mass, against 19% and 16% without it; Fig. 4c and Fig. 3c). The condition-by-readout interaction was significant (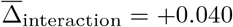, s.d. 0.067, s.e.m. 0.009; *p* = 1.9 × 10^−5^; centers of mass +0.026, *p* = 3.3 × 10^−4^). Thus, stronger field concentration during goal-directed behavior did not necessarily indicate greater parameter-dependent reorganization.

Together, these results identify distinct contributions of plasticity and teaching content. Recurrent plasticity allows the predictive spatial representation to change, while a reward-proximity teaching signal directs that change toward the rewarded location.

### Goal tuning and place-field geometry respond differently to relocation

How does the learned representation respond when the reward is moved to a new location? To address this question, we relocated the reward from location A to location B and compared changes in place-field geometry with changes in goal-directed tuning (Fig. 5). We asked how the encoding of goal-directed information and the spatial distribution of place fields change when the goal moves. To isolate changes in the learned representation from differences in behavior, we replayed the same exploratory trajectory through baseline, post-A, and post-B parameter snapshots, holding the evaluated sensory and action sequence fixed across all three conditions.

**Fig. 5.**
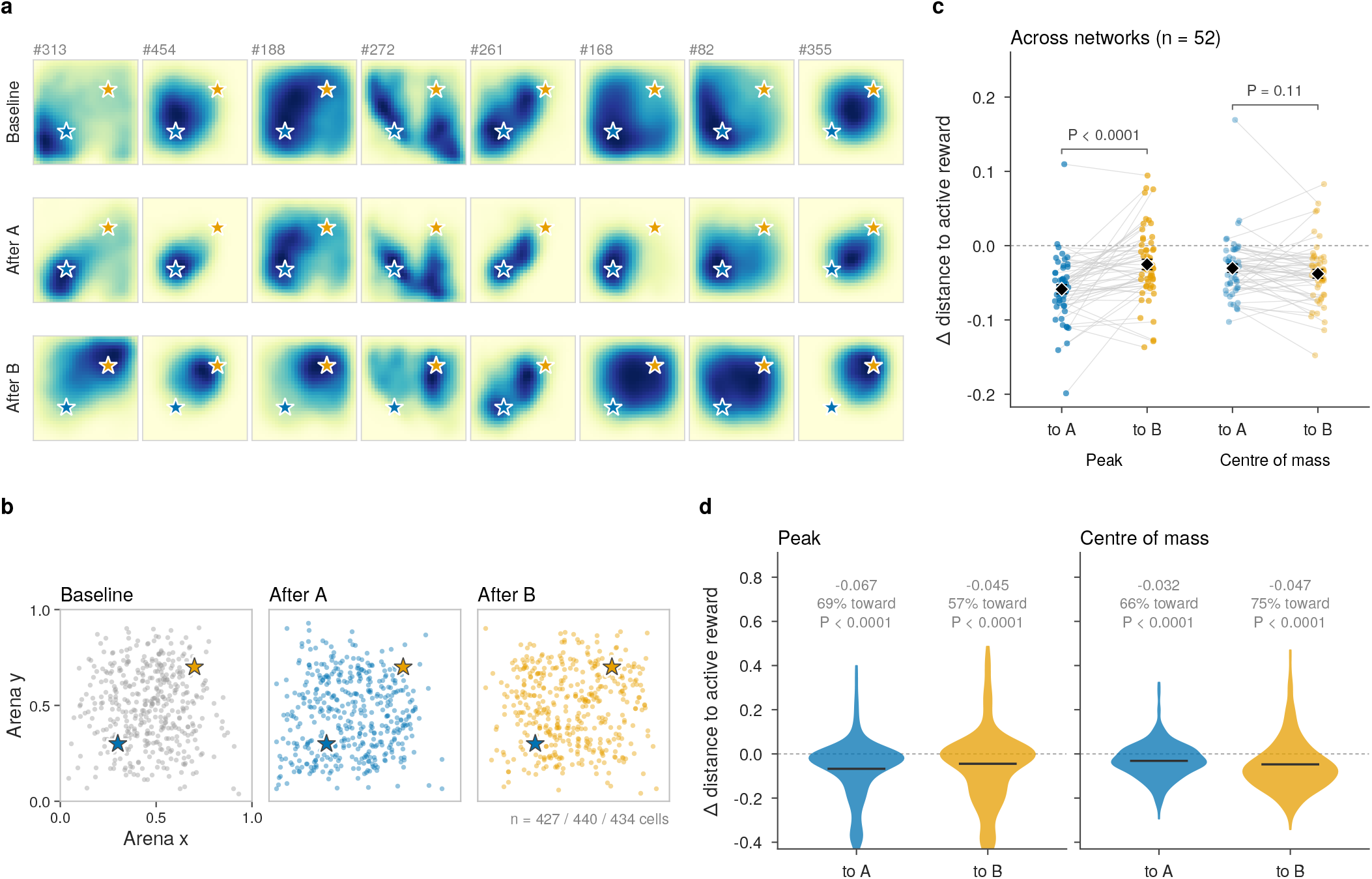
Goal-referenced bearing tuning and place-field geometry respond differently to relocation. The network first learns reward at location A, and the reward is then moved to location B. At relocation, the environmental reward, the controller’s supplied goal cue and the reward proximity target all switch from A to B. Fixed-trajectory replay passes the same stored exploratory observation–action trajectory through the network under three parameter states: baseline, after learning at reward A and after relocation to B. a) Example fields across parameter states of the network closest to the cohort median; stars, reward A (blue) and B (orange). b) Field centers of mass of the example network’s place cells in the three states (each dot is one cell; *n* = 427, 440 and 434 cells with Skaggs information ≥ 0.05 bits/spike). c) Per-network mean shift toward the active reward (arena units; negative = toward reward) for the first step (baseline to after A, toward A) and the second (after A to after B, toward B), for field peaks and centers of mass, in the 52 retained networks. Gray lines join a network’s two steps; diamonds, mean *±* s.e.m.; brackets, paired Wilcoxon tests of the difference. d) Per-cell shifts in the example network for peaks and centers of mass (same two steps as in c); horizontal lines, means; violins show the central 98% of cells, and statistics (paired Wilcoxon) use all cells; annotations give the mean shift, the percentage of cells moving toward the active reward and the P value. Across-network statistics and the recurrent-unit bearing analysis are reported in the text and Figs. S3 and S4.

We first examined the spatial distribution of place fields. In an example network, fields concentrated near A after learning and redistributed toward B after relocation (Fig. 5a,b). Across the 52 networks (Fig. 5c and Fig. S3a,b), field peaks shifted toward A after learning reward A (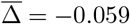, s.d. 0.044, s.e.m. 0.006; 63.0% of cells; negative in 50 of 52 networks; *p* = 5.3 ×10^−9^), as did centers of mass (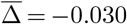, s.d. 0.041, s.e.m. 0.006; 62.3% of cells; *p* = 2.5 × 10^−7^). After relocation, fields reorganized toward B on both measures (peaks: 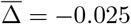, s.d. 0.050, s.e.m. 0.007; 52.7% of cells; negative in 38 of 52 networks; *p* = 5.0 × 10^−4^; centers of mass: 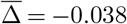, s.e.m. 0.006; 63.6% of cells; *p* = 1.2 × 10^−6^). The two measures differed in the relative size of the two steps. Field peaks moved more toward A than subsequently toward B (39 of 52 networks; paired *p* = 2.8 × 10^−5^), whereas centers of mass moved comparably toward both (*p* = 0.11; Fig. 5c and Fig. S3c). The example network showed the same pattern cell by cell (Fig. 5d). Thus, under this goal displacement, the locations of field peaks retained an influence of the previously learned reward, whereas the overall mass of the fields re-centered on the new goal about as strongly as on the first.

Goal-related measures were strongly trajectory-dependent: both the fraction of reward-distance cells and goal-bearing tuning were greater during goal-directed behavior than during replay of a random walk (Fig. S4a). We therefore used relocation, rather than tuning strength alone, to test the reference frame. On goal-directed trajectories, preferred goal bearing changed by +51.4° (95% CI [+17.4°, +88.0°]), less than the median 87° change predicted for purely allocentric tuning but also inconsistent with complete stability relative to the goal (Fig. S4b). On the fixed replay trajectory, the change was only +4.6° (95% CI [−1.3°, +10.5°]), but the allocentric prediction was also small (median magnitude 10°), so this control could not distinguish the two reference frames (Fig. S4c). The replay analysis measured the recurrent state without the goal cue supplied to the downstream controller, so that cue cannot directly account for the tuning.

Reward relocation thus affected the two measures differently. During goal-directed behavior, bearing preferences retargeted more fully than field peaks, which retained an influence of earlier learning; this asymmetry was not present for field centers of mass. This pattern suggests that directional tuning can adapt more readily than peak locations within the same recurrent population, but it does not establish a fully goal-centered code. Place-cell vector fields and hippocampal theta sweeps that represent goal direction independently of movement and heading provide a biological parallel (32, 33). Our results do not establish how such signals arise in the brain, how quickly they update, or whether the model and hippocampal theta sequences use the same mechanism.

### Withdrawing reward and proximity teaching reverses field concentration near reward

Does the spatial organization acquired during reward learning persist when the signals that established it are withdrawn? To connect this question to recordings of CA1 place cells and VTA axons during extinction of reward expectation (34), we compared the experimental protocol with a controlled model manipulation (Fig. 6a). After rewarded learning, we removed both environmental reward and the proximity-objective gradient, while retaining the supplied goal cue and goal-triggered termination, and replayed the same exploratory trajectory through the baseline, learned and withdrawn networks. This protocol does not model how an animal infers that reward has disappeared instead, it asks whether withdrawing two specified learning signals is sufficient to reverse the parameter-dependent organization they produced.

**Fig. 6.**
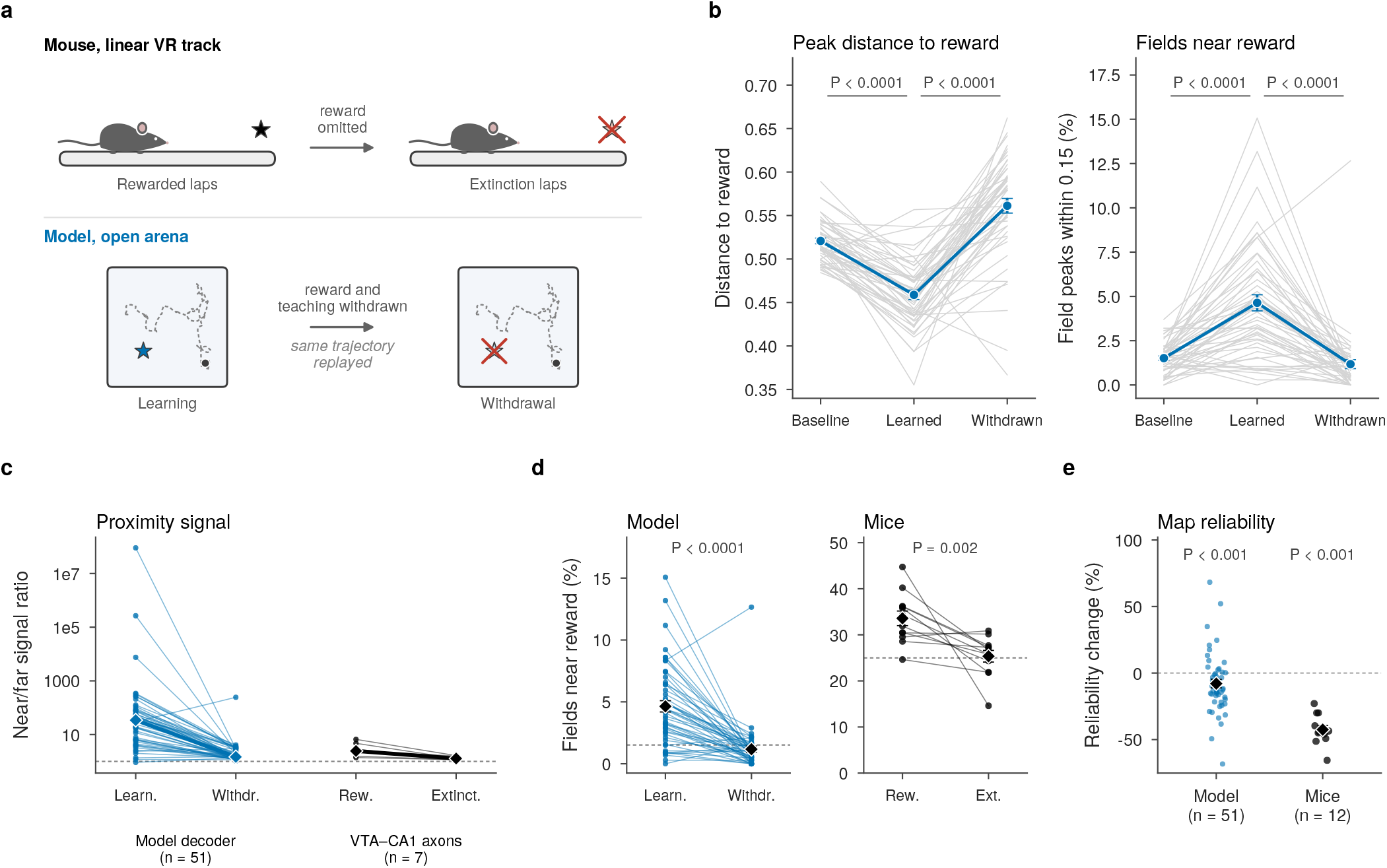
Limited correspondence with a reward-omission experiment. After reward learning with proximity teaching, environmental reward and the proximity teaching signal are withdrawn together, while the supplied goal cue and goal-triggered termination remain. Fixed-trajectory replay passes the same stored exploratory observation– action trajectory through the network under three parameter states: baseline, learned and withdrawn. Model quantities are blue and experimental quantities black; diamonds and error bars show mean *±* s.e.m. unless stated otherwise. a) Protocols: mice on a linear virtual-reality track (top) and the model in an open arena (bottom). b) Model (*n* = 51 networks; gray lines, individual networks; place cells with Skaggs information 0.15 bits/spike in all three states): mean distance of field peaks to the former reward (arena units; left) and percentage of field peaks within 0.15 arena units of it (right). Learning concentrates fields toward the reward and withdrawal reverses the change with overshoot (recovery fraction 1.65, where 1 is a return to baseline; bootstrap 95% CI [1.37, 1.94]); brackets, paired Wilcoxon tests. c) Proximity signal: near/far output ratio of the model auxiliary readout (weights fixed after learning; *n* = 51) and end/start activity ratio of VTA–CA1 axons in rewarded and low-expectation extinction laps (Fig. 5d of Krishnan et al.; *n* = 7 axons); lines join a network or axon, thick lines join medians. The model readout has a much steeper target than the measured ramp. d) Fields near the reward before and after withdrawal: model field peaks within ≥0.15 arena units of the former reward (dashed line, baseline mean) and recorded fields in the reward region (dashed line, 25% expected from a uniform distribution), paired per network or animal (*n* = 51 networks, 12 animals). e) Change in split-half map reliability from learned to withdrawn (model, per network) and from rewarded to extinction laps (mice, per animal; *n* = 12, Supplementary Fig. 11b of Krishnan et al.). These comparisons are descriptive, not fitted.

Withdrawal was sufficient to reverse the learned concentration of fields near reward. Learning moved field peaks toward the reward (mean change in distance −0.062, s.d. 0.039, s.e.m. 0.005; *p* = 1.3 × 10^−9^, one-sided), whereas withdrawal moved them away by more (+0.102, s.d. 0.070, s.e.m. 0.010; *p* = 3.5 × 10^−9^, one-sided). The endpoint passed the original baseline (recovery fraction 1.65, bootstrap 95% CI [1.37, 1.94]), and the fraction of peaks near reward rose from 1.5% to 4.6% during learning before falling to 1.2% after withdrawal (Fig. 6b). Thus, reward-related organization was not frozen into the predictive map: continued learning without the two reward-related signals actively reorganized it. The overshoot also shows that the implemented dynamics do not simply restore the baseline map and should not be interpreted as a quantitative fit to extinction.

This representational reversal was visible to a downstream readout without changing the readout itself. The proximity decoder received no updates during withdrawal, yet its median near-to-far output ratio fell from 34 to 1.5 (*p* = 1.1 × 10^−8^; Fig. 6c and Fig. S5a). VTA–CA1 axons showed a qualitatively parallel flattening, with a median end/start ratio of 2.4 → 1.3 across rewarded and low-expectation laps (Fig. 6c). The much steeper initial contrast in the model rules out treating its decoder as a quantitative match to the measured axonal signal, the stronger mechanistic conclusion is that changing the recurrent representation alone can substantially weaken the proximity signal expressed by an established readout.

The place-field comparison provides a second parallel. In the model, near-reward peaks returned from their learned enrichment to approximately baseline, while in mice the fraction of fields in the reward region fell from 33.6% to 25.4% (paired rewarded-versus-low-expectation *p* = 0.002), with the latter close to the 25% expected by chance (Fig. 6d). These measures are not directly commensurate, but both show that an acquired concentration near reward is lost after withdrawal.

The comparison also identifies what withdrawal of the implemented signals cannot explain. Map reliability declined in both systems, but much less in the model than in mice (−7.9% per network versus −42.5% per animal, s.e.m. 3.2 for both; Fig. 6e). Moreover, the model’s decline was confined to cells far from the former reward, whereas the experimental decline extended across track regions (Fig. S5b). Behavior directed at the former reward nevertheless stopped rapidly in both systems within tens of episodes in the model and a few laps in mice (Fig. S5c,d). Together, these comparisons show how the model can be used to test which experimental effects are captured by its simple mechanisms and which require additional explanations.

### CA1 reward-relocation data show partial correspondence with the model

Do hippocampal maps respond to reward relocation as the model does? We reanalyzed CA1 recordings from mice on a virtual track in which the reward zone moved within a session, with anticipatory licking recorded concurrently (seven switch sessions from two mice; DANDI 001361; Fig. S6a) (35), and applied to them the measures used for the model’s relocation from A to B (Fig. 7a; Methods).

**Fig. 7.**
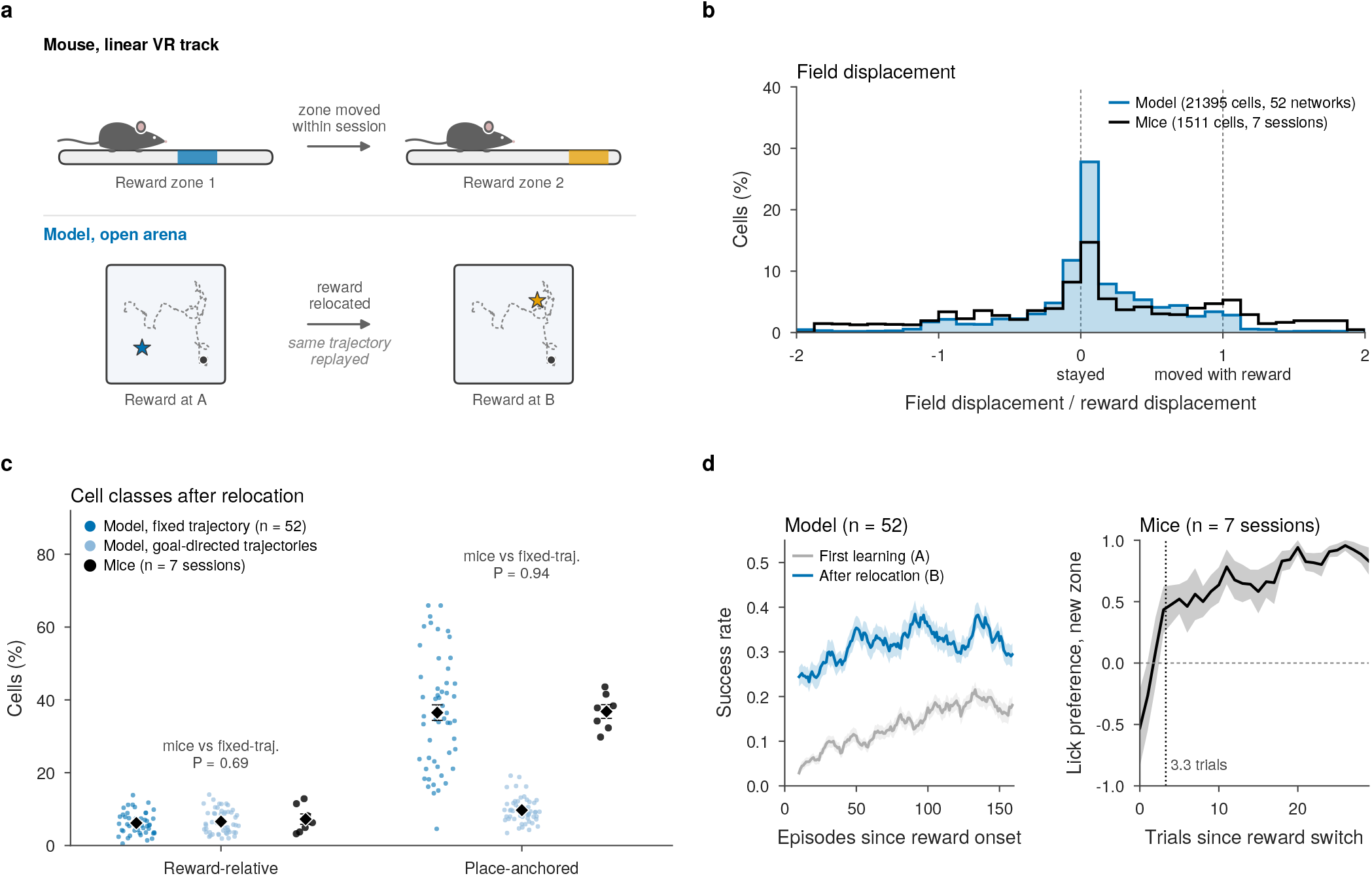
Limited correspondence with a reward-relocation experiment. The reward is moved within a session, from zone 1 to zone 2 for mice (Sosa et al.) and from A to B for the model, where the goal-displacement cue and the reward proximity target switch together. Unless stated otherwise, model networks are read out with fixed-trajectory replay of the same stored exploratory observation–action trajectory before and after relocation. Model quantities are blue and experimental quantities black; diamonds and error bars show mean *±* s.e.m. a) Protocols: mice on a virtual-reality track (top) and the model in an open arena (bottom). b) Displacement of each field peak across the switch, in units of the reward displacement (0, field stayed; 1, field moved with the reward): model cells pooled over networks (*n* = 21,395 cells, 52 networks; fixed-trajectory replay, displacement projected on the A–B axis) and recorded place cells pooled over sessions (*n* = 1,511 cells, 7 sessions; displacement on the track). Within 0.2 of 0: 48% of model and 29% of recorded fields; within 0.2 of 1: 8% and 14%. c) Reward-relative and place-anchored cell fractions under the same classification, for model networks (*n* = 52) read out with fixed-trajectory replay (dark blue) and on goal-directed trajectories (light blue), and for recording sessions (*n* = 7, from two mice; black). P values are one-sample Wilcoxon tests of the sessions against the fixed-trajectory model mean; they detect no difference but are not evidence of equivalence. d) Behavioral retargeting. Left, model success rate per episode after relocation to B and during initial learning of A (10-episode running mean; mean *±* s.e.m. over 52 networks); the controller receives the displacement to the current goal. Right, mouse anticipatory-lick preference for the new zone, (new − old)/(new + old), after the switch (3-trial running mean; mean *±* s.e.m. over 7 sessions; dotted line, mean lick latency).

When the reward moved, fields in both systems split in the same way: most stayed near their previous location, and a smaller group moved by about the displacement of the reward (Fig. 7b; an example session in Fig. S6b). The group that stayed was more sharply defined in the model.

Applying the same classification rule to both systems yielded similar cell-class fractions when the model was evaluated along a repeated trajectory. Place-anchored cells, whose fields remained in place, comprised 36.8% of recorded cells (s.e.m. 1.9) and 36.5% of model units (s.e.m. 2.1); reward-relative cells, whose fields followed the reward, comprised 7.2% (s.e.m. 1.4) and 6.2% (s.e.m. 0.4), respectively (data: seven sessions; model: 52 networks; Fig. 7c). By contrast, evaluating the same model networks along goal-directed trajectories reduced the place-anchored fraction to 9.7%, showing that cell classification depends on the sampled trajectory. Because the mice followed nearly the same route on successive laps, the repeated-trajectory model readout provides the more appropriate comparison. Nevertheless, this numerical similarity is not evidence of equivalence: the estimated fractions depend on the classification rule (Fig. S6c), and the experimental sample comprises only seven sessions. Behavior switched to the new reward almost at once in both systems (Fig. 7d). Model success at B was 0.24 in the first ten episodes after relocation, against 0.03 at the start of initial learning, and mice shifted anticipatory licking to the new zone within 3.3 trials (s.d. 1.9). In the model this speed has a simple origin: the new goal reaches the controller directly, whereas the map changes only through plasticity, so behavior can switch before the map does. The recordings are consistent with this ordering but do not establish it: the density of fields near the new zone was estimated to transfer after 11.0 trials, later than licking in five of six decidable sessions (*p* = 0.0625), but the estimate cannot resolve transfers faster than five trials (Fig. S6d,e).

Together, the reanalysis shows two descriptive parallels with the model: comparable cell-class fractions under one shared rule, and behavior that adjusts before the place-field map. Given the two-animal sample and the five-trial floor on the map estimate, both findings need broader replication, and showing that the responses arise from the model’s mechanisms will require direct perturbation.

### Behavioral matching reduces sampling effects but does not isolate map reorganization

Can the changes isolated by fixed-trajectory replay be recovered from recordings made during freely varying behavior? The repeated track trajectories in the experimental reanalysis provide a relatively constrained setting: there, matching occupancy barely changed the estimated field shift (*p* = 0.94, seven sessions; Fig. S6f). Exploration and goal-directed navigation in the open arena, by contrast, produce different paths and headings. We therefore matched the joint distribution of position and heading between pre-learning exploration and post-learning goal-directed behavior, then compared the resulting field shifts with fixed-trajectory replay. The analysis used 55 matched network pairs with and without proximity teaching and examined activity both before and after the interleaved sensory-prediction training (Methods; Fig. S7).

Matching substantially reduced the apparent concentration of fields near reward. Before interleaved training, unmatched rate maps gave mean shifts of 0.208 (s.d. −0.040, s.e.m. 0.005) without proximity teaching and −0.191 (s.d. 0.036, s.e.m. 0.005) with it, compared with −0.039 (s.d. 0.053, s.e.m. 0.007) and −0.062 (s.d. 0.032, s.e.m. 0.004) under fixed-trajectory replay. Matching position and heading reduced the behavioral estimates to −0.043 (s.d. 0.024, s.e.m. 0.003) and −0.027 (s.d. 0.019, s.e.m. 0.003), respectively (Fig. S7a). A control using identical network parameters for both trajectories confirmed that much of the unmatched shift arose from sampling: apparent shifts of −0.148 (s.d. 0.059, s.e.m. 0.008) and −0.121 (s.d. 0.045, s.e.m. 0.006) fell to −0.007 (s.d. 0.029, s.e.m. 0.004; *p* = 0.036) and +0.002 (s.d. 0.021, s.e.m. 0.003; *p* = 0.36) after matching (Fig. S7b). Thus, matching removed most of the apparent shift in this control, although a small residual remained without proximity teaching.

Despite this improvement, matching did not recover the effect of proximity teaching. Agreement with fixed-trajectory replay across individual networks was weak (*r* = −0.06 without teaching and *r* = 0.21 with it; Fig. S7c). Moreover, whereas replay showed a larger shift toward reward with the teaching objective (*p* = 1.8 × 10^−3^), matched behavioral recordings gave the opposite ordering (−0.043 without teaching versus −0.027 with teaching; *p* = 4.6 × 10^−4^; Fig. S7d). The ordering remained reversed after interleaved predictive training, and agreement with fixed-trajectory estimates remained weak.

Matching current position and heading is not enough for a recurrent network, because its activity also depends on the sensory and action history leading to that state. The matching procedure removed much of the average sampling bias, but it did not recover the size of the parameter-dependent shifts or their ordering across conditions. Thus, within the model, controlling the full evaluated sequence remains necessary to separate changes in sampling from changes caused by learning.

## Discussion

Reward-related changes in spatial activity can arise because behavior alters how an existing representation is sampled or because learning changes the representation itself. We separated these contributions in a recurrent network that acquired spatial tuning through sensory prediction before exposure to reward. Goal-directed behavior strongly concentrated measured fields near reward, whereas replaying the same sensory–action sequence before and after learning revealed a smaller change attributable to the learned parameters. This parameter-dependent reorganization became larger and more consistently directed toward reward when learning included an explicit proximity objective, and was attenuated when recurrent plasticity was restricted. Relocation and withdrawal further showed that reward-related features of the representation could update or reverse without uniformly destabilizing the spatial map. Together, these findings distinguish the behavioral expression of a predictive spatial representation from its modification by reward-related learning.

### Comparison with hippocampal reward coding

Without being fitted to any one experiment, the model links several qualitative findings usually studied separately. Sensory prediction generates place-cell-like tuning before reward, and this spatial map remains organized during goal learning while fields become concentrated and shift toward reward (5, 6, 11– 13, 15, 23–26). Reward-proximity teaching turns variable plasticity into a reliable reward-directed change, consistent with a role for catecholaminergic modulation and VTA–CA1 proximity signals (8, 10, 34, 36–38). These correspondences are informative without implying quantitative equivalence.

Relocation produced both place-anchored and reward-relative responses and revealed different degrees of flexibility within the same population: field peaks retained an influence of the first reward, whereas preferred bearings partially retargeted toward the current goal (35). This pattern is consistent with coexisting space- and goal-referenced inputs whose balance can be altered by experience and behavioral-timescale synaptic plasticity (39). It does not, however, establish a fully goal-centered code: the goal-directed response was intermediate between complete goal stability and the allocentric prediction, the replay control was geometrically ambiguous, and the model lacks the within-theta-cycle dynamics of goal-oriented sweeps (16, 32, 33, 40). The CA1 reanalysis provided a further descriptive parallel, yielding similar cell-class fractions under a shared classifier and suggesting that behavior may adjust before the field distribution. These observations require broader testing because the dataset comprised seven sessions from two animals and the map analysis could not resolve changes within five trials (35).

Reward withdrawal identified a second boundary of the model. Near-reward enrichment and proximity contrast declined in both model and data, but the model began with a much steeper decoder contrast and showed a smaller, spatially restricted loss of map reliability rather than the widespread destabilization observed experimentally (34). The manipulation also removed environmental reward and proximity teaching together while preserving the goal cue and goal-triggered termination. It therefore tests the sufficiency of withdrawing specified learning signals, not how an animal infers that reward is no longer available. Recent evidence linking hippocampal activity to latent-state inference suggests one mechanism that may contribute to this missing biological response (41).

### Separating sampling from representational change

Interpreting these correspondences requires separating changes in how a representation is sampled from changes in the representation itself. A rate map averages activity across visits to the same location, but the state of a recurrent network also depends on the path and sensory history leading there. A behavioral change can therefore reshape a measured field without changing the network parameters, particularly near reward, where trajectories, headings and recent inputs differ from exploration. In our model, this sampling effect was large enough to reverse the apparent ordering of the proximity-teaching conditions. Field concentration during behavior is therefore not a direct measure of representational change.

Fixed-trajectory replay isolates the parameter-dependent component by holding the initial state and sensory–action sequence constant. It measures change along the tested trajectory, not a trajectory-independent property, and does not simulate hippocampal replay. Matching position and heading reduced average sampling bias but left sensory history uncontrolled and failed to recover the ordering of the learning conditions. Repeated routes and tighter control of recent experience are therefore useful experimental approximations, although they cannot match the control available in the model.

Geometry creates a second confound. A localized field can correlate with distance to any fixed point before that point has behavioral significance, and regression toward the arena center can make field displacement appear reward-directed. Requiring activity to be uniform around iso-distance rings greatly reduced apparent reward-distance coding, and the remaining distance and bearing measures depended strongly on the sampled trajectories (Fig. S4a). The relocation comparisons further show why an explicit reference-frame control is needed to interpret goal-bearing tuning (Fig. S4b,c).

### Reward-related teaching in a predictive spatial representation

Sensory prediction provides a route to spatial organization before reward is introduced (26). In the model, continued prediction preserved spatial tuning, while proximity teaching biased where fields were expressed along a fixed trajectory. Recurrent plasticity permitted fields to reorganize, whereas teaching content directed more of that change toward reward without increasing total displacement. Thus, a predictive map can retain its overall organization while being shaped by behavioral demands (6). The tested plasticity settings support this dissociation, but do not establish a monotonic relationship between learning rate and reward-directed change.

In its standard form, the successor representation contains no reward or goal signal: it represents expected future state occupancy under the current policy, with reward handled by a separate readout (19–21). It changes here only because reward learning changes the policy and therefore the agent’s trajectories. This policy-dependent change explains part of the behavioral field shift (Fig. S8a), but not the residual change under fixed-trajectory replay: for an identical trajectory the successor representation predicts no change, whereas the recurrent fields still shift toward reward (Fig. S8b). The recurrent model therefore captures an additional parameter-dependent change beyond policy-dependent occupancy.

Two assumptions constrain the biological interpretation. First, the controller is given the true displacement to the goal. Orbitofrontal goal representations and hippocampal sequences and vector fields directed toward remembered or current goals show that goal information is available in the brain, but do not specify the source or coordinate system assumed by the model (16, 29–33, 42). Second, the recurrent network is taught a smooth function of true reward distance. Dopaminergic modulation of hippocampal plasticity and VTA–CA1 reward-proximity ramps, distinct from LC novelty and movement signals, make such an instructive pathway plausible (34, 36–38). The supplied target is nevertheless neither a model of dopamine activity nor a reward-prediction error. Because the controller’s reward already includes progress toward the goal, the comparison isolates the added effect of explicit proximity supervision rather than learning from sparse reward alone.

### Experimental predictions and limitations

Prior VTA and LC manipulations show that catecholaminergic inputs can direct or suppress reward-associated place-field redistribution (8, 10). The model makes the stronger prediction that suppressing a reward-related teaching input should reduce the acquired concentration of fields near reward more than it impairs navigation. Comparing matched perturbations of VTA and LC inputs during repeated routes, together with hippocampal and behavioral measurements, could determine whether this dissociation is specific to reward proximity rather than novelty or movement signals.

Relocation and omission offer complementary tests of how representations update over time. During relocation, trial-resolved measurements of dendritic plateau events, spatial and goal reference frames, field organization and behavior could establish whether these signals change at different rates; our endpoint comparisons cannot. During omission, a decoder trained before withdrawal and then held fixed, together with a latent-state model of behavior, could distinguish loss of a teaching signal from inference of a new context. Refitting the decoder would further separate a change in coding format from a loss of decodable information. Trial-resolved recordings could also test for the backward progression of reward-related activity toward earlier predictive states (27).

The conclusions remain limited to the tested environments, reward locations, plasticity settings and learning regimes, and should be evaluated across a broader range of conditions and contexts (9). A key extension is to replace the supplied goal and proximity variables with signals learned from sensory history and reward outcomes. This would test whether the same separation persists when the system must discover goals, infer reward availability and construct its own teaching signal.

Predictive spatial representations can therefore guide goal-directed behavior while remaining open to reward-dependent modification. Separating how a map is sampled from how it is changed provides a framework for determining when reward-related inputs guide behavior, reshape spatial representations or do both.

## Acknowledgments

We would like to thank Omri Barak and the many people with whom we had fruitful discussions about the development of this work. In particular, we thank Rich Pang for inspiring conversations about world models and non-Hebbian plasticity that informed the predictive component of this model.

## Data and materials availability

Code is available at https://github.com/TechnionNeuroAILab/spatial_predictive_RL. The recordings reanalyzed here are available on the DANDI Archive (dandiset 001361).

## Methods

The methods follow the causal sequence of the experiments. We first describe the arena, sensory input and recurrent network, we then describe predictive pretraining (Phase A), goal-directed reinforcement learning (Phase B), and the optional reward-proximity objective, finally, we define the replay controls, inclusion criteria and spatial statistics. Three readouts recur throughout the paper. The *pre-learning exploratory readout* uses a random-walk trajectory and the Phase A parameters. The *post-learning behavioral readout* uses a goal-directed trajectory and the Phase B parameters, so both parameters and sampled experience differ. The *fixed-trajectory readout* passes one stored random-walk sequence through the pre- and post-learning parameters, so only the parameters differ. This distinction determines whether a reported change describes activity expressed during behavior or a change in the recurrent representation under matched input.

### Training algorithm

The full pipeline comprises predictive training, cyclic reinforcement learning with interleaved predictive training, the optional reward-proximity decoder and a fixed-trajectory replay probe. Algorithm 1 gives the chronology; the sections that follow define each component and analysis in detail.

Here *E*_1_ denotes the number of Phase A epochs, *N*_ep_ the total number of Phase B episodes and *N*_cyc_ the number of Phase B cycles, the designated replay cycle was the first cycle in all replay analyses. Numerical values are collected in Table S1.

### Environment

We used the RatInABox framework (43) to simulate a 2D bounded arena (1 × 1 unit square). Positions and Euclidean distances are therefore reported in arena units. Objects of three types (12 per type) were placed at random locations. The agent navigated on a 32 × 32 grid with 8 possible movement directions (cardinal + diagonal). Requested continuous reward coordinates were converted to the nearest grid point for the training episode’s goal cue and termination condition: (0.3, 0.3) became (10*/*32, 10*/*32) = (0.3125, 0.3125) and (0.7, 0.7) became (22*/*32, 22*/*32) = (0.6875, 0.6875). The proximity target, goal-directed probes and spatial analyses instead used the requested continuous coordinates. Phase A training trajectories followed a heading-biased random walk: relative to the current heading, the agent moved straight ahead with probability 0.5, turned by ±45° with 0.15 each, by ±90° with 0.05 each and by ±135° with 0.04 each, and reversed with 0.02; moves into a wall were removed and the remaining probabilities renormalized. The 50,000-step random-walk probes used for place-field analysis and fixed-trajectory replay drew each of the eight moves with equal probability, clipped at the walls. Goal-directed probes recorded after each RL cycle contained 5000 recorded steps. A separate final post-interleaving goal-directed probe used for tuning analyses contained 50,000 recorded steps. Each probe comprised 50 random-start segments, each beginning with 10 unrecorded random warm-up moves, and also reset the agent and recurrent state when it came within 0.06 of the requested reward coordinate (training used the 0.05 radius in Eq. (S9) around the snapped coordinate), the reset handling used in the rate maps is defined below.

### Observations

At each timestep *t*, the agent at position ***x***_*t*_ with heading *θ*_*t*_ received egocentric observations ***o***_*t*_ ∈ ℝ^*M*^ consisting of:

- **Object-sensitive Field-of-View cells**: three populations of six RatInABox field-of-view object-vector cells (43), one population per object type, with pre-ferred egocentric angles *ϑ*_*i*_ ∈{± 8°, ±24°,± 40°}. Cell *i* of type *k* sums over the objects *o* of that type at positions ***p***_*o*_:

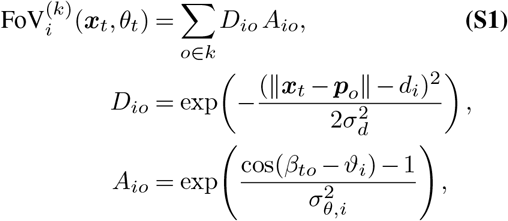

with distance tuning *D*_*io*_ and angular tuning *A*_*io*_, where *β*_*to*_ is the egocentric bearing of object *o* relative to heading *θ*_*t*_ and *σ*_*d*_ = 0.05. The preferred distance *d*_*i*_ = 0.99 *w*_*t,i*_ and angular width *σ*_*θ,i*_ = 0.0505*/w*_*t,i*_ (radians) scale with *w*_*t,i*_, the distance to the nearest wall along the cell’s preferred direction.
- **Wall distance rays**: distances to the nearest wall along each tuning angle, normalized by 1.5.

Total observation dimensionality was *M* = 24: three object types × six egocentric field-of-view cells (firing rates, scaled by 1*/*10) concatenated with six ray-cast wall distances (scaled by 1*/*1.5).

### RNN architecture

The predictive RNN had hidden dimension *H* = 512 and received the current observation ***o***_*t*_ ∈ ℝ^*M*^ and an eight-dimensional one-hot action code ***ã***_*t*_. Its hidden state evolves as

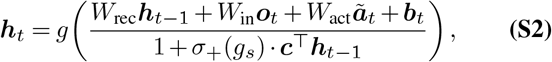

with a sparse, biased-ReLU activation

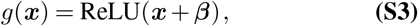

where ***β*** ∈ ℝ^*H*^ is a learned bias initialized to −0.2; the negative bias produces exact zeros for sub-threshold inputs, yielding the sparse, place-cell-like coding seen in Fig. 2. The denominator implements divisive inhibition through a learned readout ***c*** and a gain *g*_*s*_ passed through a softplus *σ*_+_(*x*) = log(1 + *e*^*x*^) to ensure positivity. The bias ***b***_*t*_ is a slow memory trace that integrates observation history,

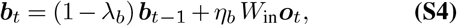

with *λ*_*b*_ = 0.1 and step size *η*_*b*_ = 0.1 (so the trace retains a fraction 1 − *λ*_*b*_ = 0.9 of its previous value each step). The output layer predicts the next observation via a sigmoid, 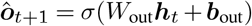.

#### Algorithm 1

Predictive RNN, goal-conditioned controller and reward-proximity objective

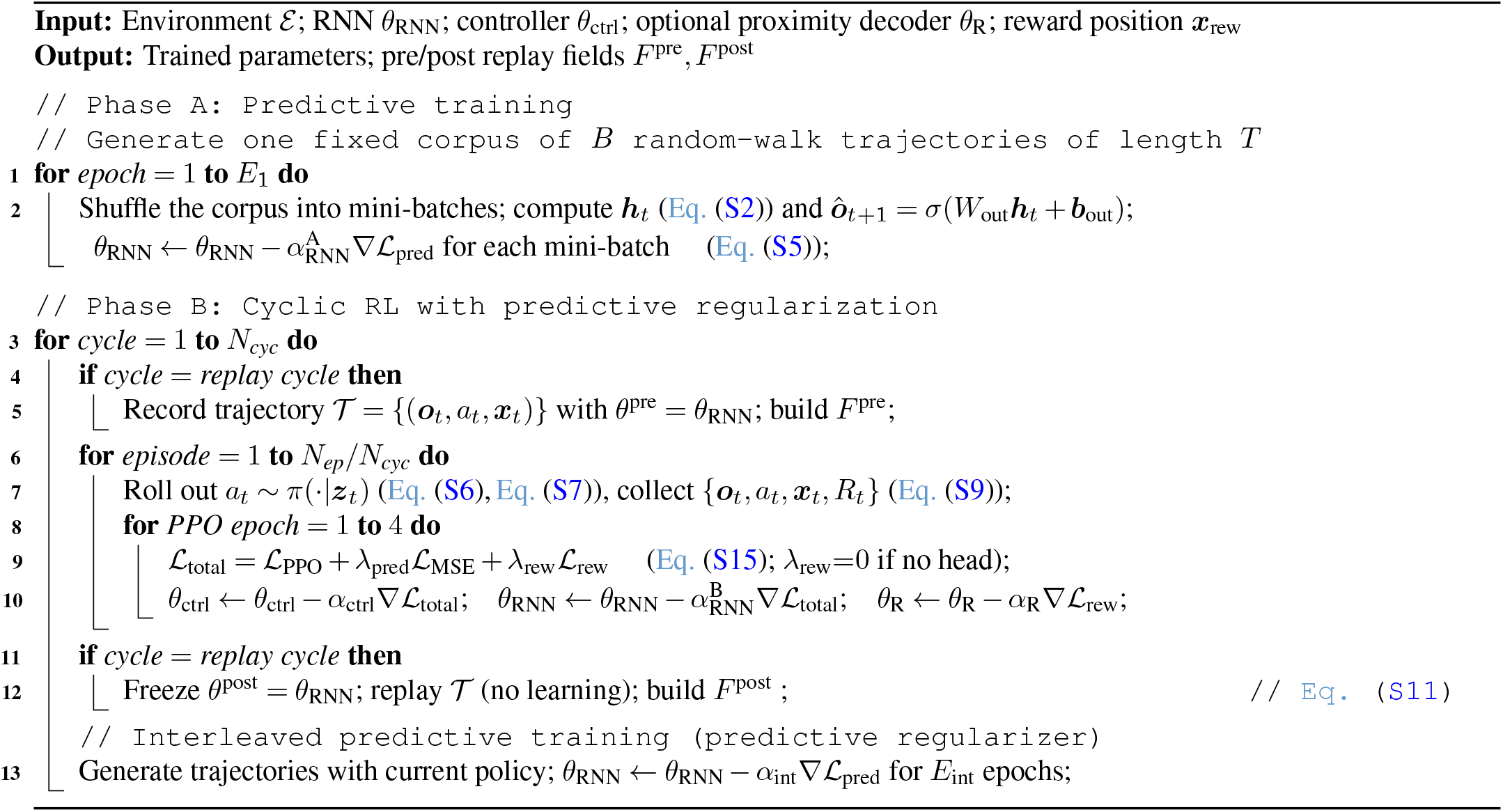

Total learnable parameters in Phase A: *W*_rec_ ∈ ℝ^*H×H*^ , *W*_in_ ∈ ℝ^*H×M*^ , *W*_act_ ∈ ℝ^*H×*8^, *W*_out_ ∈ ℝ^*M ×H*^ , ***b***_out_ ∈ ℝ^*M*^ , ***c*** ∈ ℝ^*H*^ , *g*_*s*_ ∈ℝ, ***β*** ∈ ℝ^*H*^. In Phase A and the interleaved predictive batches, ***ã***_*t*_ is the outgoing action that generates ***o***_*t*+1_. During online control and all analysis probes, the causal state update instead receives the action on the preceding transition (equivalently, the current movement-heading code), the within-episode Phase B prediction regularizer retains this one-step-lagged action convention while predicting the next observation. We preserve and report this distinction because the released cohorts were generated with it.

### Phase A: Predictive training

The RNN was trained on one corpus of *B* = 1000 trajectories of *T* = 100 recorded timesteps each, generated by the heading-biased random walk described above. The adjacent-step construction supplied the first 99 observation–action pairs as inputs and the following 99 observations as targets. The same corpus was shuffled into mini-batches on each epoch, trajectories were not regenerated between epochs. Training minimized the next-step prediction loss augmented with L1 activity regularization,

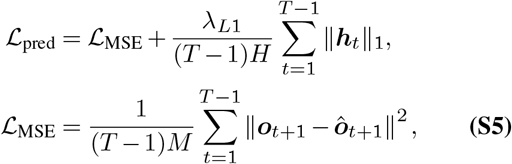

averaged over the mini-batch, with *λ*_*L*1_ = 10^−4^, over 100 epochs with RMSProp (lr = 0.001), batch size 64. Weights were initialized uniformly on 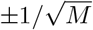 for *W*_in_ and *W*_act_ and on 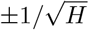 for *W*_rec_ and *W*_out_, with the recurrent matrix additionally given a leaky-integrator identity component, *W*_rec_ ← *W*_rec_ + (1 − 1*/τ*) *I* with *τ* = 2. Gradients were clipped to norm 1.0 in Phase A, in Phase B to 1.0 (RNN), 0.5 (controller) and 0.5 (proximity decoder), and to 0.5 during interleaved predictive training; Adam used *ϵ* = 10^−5^.

### Goal-conditioned controller architecture

The controller received the RNN hidden state ***h***_*t*_ and an explicitly supplied allocentric goal-displacement vector ***g***_*t*_ = (*g*_*x*_ − *x*_*t*_, *g*_*y*_ − *y*_*t*_). This is the Cartesian displacement in arena coordinates, not the direction relative to the agent’s heading. It was computed from the true agent and goal positions at every step and entered the controller additively: with the layer-normalized hidden state 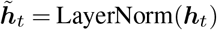,

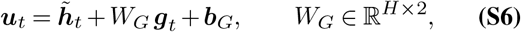

a learned affine transformation of the goal displacement added to the recurrent state. After a two-layer residual MLP, ***z***_*t*_ = ***u***_*t*_ +MLP(***u***_*t*_), separate actor and critic heads produce

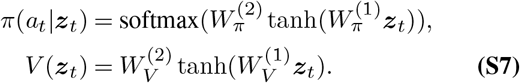

In the released code *W*_*G*_ and ***b***_*G*_ are parameterized as a goal-encoder layer followed by the value and output projections of a single-token attention module with a single key the attention weights are identically one, so this composition reduces exactly to the affine map in Eq. (S6). The agent nevertheless has a movement direction: actions select one of eight allocentric movements, and the egocentric sensory observation depends on the resulting heading. Heading is not concatenated directly with the goal displacement. Concrete layer dimensions:

- Additive goal input: affine map ℝ^2^ → ℝ^*H*^ (*W*_*G*_, ***b***_*G*_), added to LayerNorm(***h***_*t*_)
- MLP: Linear(*H* → *H*) → ReLU → Linear(*H* → *H*)
- Actor head: Linear(*H* → 64) → Tanh → Linear(64 →8)
- Critic head: Linear(*H* → 64) → Tanh → Linear(64 →1)

### Phase B: Cyclic RL training and interleaved predictive training

The controller was trained with Proximal Policy Optimization (PPO) (44) using the clipped surrogate objective

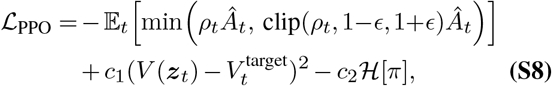

where *ρ* = *π*(*a* | ***z***_*t*_)*/π* (*a* | ***z***_*t*_) is the probability ratio and *Â*_*t*_ the advantage, estimated as a discounted return minus the critic baseline and normalized per episode. Returns were bootstrapped from the critic when an episode reached the step limit without reward, reward-terminated episodes used zero terminal value. Each training episode sampled an interior start at least five grid-index units from the goal, initialized the recurrent hidden and slow-bias states to zero, and then took 10 uniformly random, unoptimized warm-up moves before the PPO rollout. Rollouts with fewer than two recorded steps were retained in the behavioral history but skipped for parameter updates. During each update epoch, recurrent states were regenerated from the stored episode sequence and gradients did not cross episode boundaries. The main and relocation simulators initialized this recomputation from the detached pre-rollout hidden and slow-bias states. The omission simulator retained an earlier indexing convention: it initialized from the first stored post-step states and then processed the first stored observation–action pair again. This one-step exception is preserved in the released code to reproduce the reported omission cohort; all omission contrasts remain within the same networks and implementation. The environment reward combined a goal bonus, progress shaping and a step penalty,

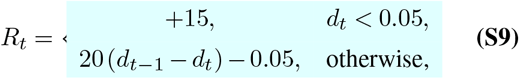

with *d*_*t*_ = *d*(***x***_*t*_, ***x***_goal_) and episodes truncated at *T*_max_ = 20 steps. Gradients from the RL loss were allowed to flow into the RNN at a reduced learning rate, alongside the mean-squared next-observation term from Eq. (S5),

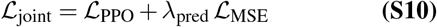

(or Eq. (S15) when the proximity decoder was used). The online PPO update did not include the L1 term. The full prediction loss, including L1 activity regularization, was restored in the interleaved predictive block. The task reward is itself dense and spatially informed: in addition to a terminal bonus it contains signed progress toward the true goal. Thus the comparison tests whether an explicit state-level proximity objective changes the representation beyond an already shaped policy/value objective, it is not a comparison with learning from sparse biological reward alone. Other hyperparameters were *γ* = 0.99, clip *ϵ* = 0.1, four PPO update epochs per episode, entropy coefficient *c*_2_ = 0.02 and value-loss coefficient *c*_1_ = 0.5. RL was organized into cycles for the replay experiments (Fig. 3, Fig. 4) the goal was fixed for one Phase B cycle of 1000 episodes. The controller used *α*_ctrl_ = 10^−4^ and the RNN 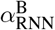 (between 10^−6^ and 10^−4^; see below), as parameter groups in one Adam optimizer.

Each cycle followed the same order: the replay-source sequence was recorded at the current weights in the designated replay cycle, RL episodes were then completed. The stored sequence was replayed through the post-RL weights, a 10,000-step unbiased random-walk probe and a 5000-step goal-directed probe were recorded and interleaved predictive training updated the RNN on the full ℒ_pred_ (Eq. (S5)) for *E*_int_ = 20 epochs on 1000 trajectories with the same 100-record/99-transition construction as Phase A, generated greedily by the current policy without early termination (a fresh Adam optimizer each cycle, lr 10^−4^). Training rollouts sampled the untempered policy, goal-directed analysis probes sampled after dividing the policy logits by 0.3. Thus, the fixed-trajectory “post-RL” snapshot and the post-RL behavioral probes precede interleaved training. The primary replay contrast isolates parameter changes produced during joint PPO/MSE training within the RL block, analyses that use the later post-interleaving snapshot are identified explicitly.

### Fixed-trajectory replay control

To dissociate parameter-dependent activity change from trajectory and occupancy, before RL we ran a random-walk probe of length *T*_replay_ = 50,000 and saved its complete observation/action sequence and positions, 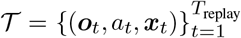. We evaluated this same stored sequence without learning through the pre- and post-RL parameter snapshots, producing *F* ^pre^ and *F* ^post^. Because observations, actions, positions and initial states are identical, any difference is attributable to the parameter snapshot:

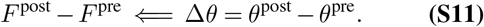

Both passes initialize ***h***_0_ = ***b***_0_ = **0**. For each cell *j* we measured the change in field-center distance to reward,

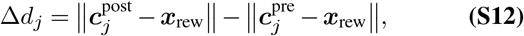

where Δ*d*_*j*_ *<* 0 indicates a field that moved toward reward under identical sampling. Paired-cell inclusion criteria are defined below under Spatial analysis and shift metrics.

This is a computational control, not a model of hippocampal replay: the stored sequence is imposed offline only to equalize input and occupancy. It estimates parameter-dependent activity change conditional on that trajectory and should not be interpreted as a trajectory-independent distance between maps.

### Reward-proximity decoder and teaching objective

The optional decoder is a two-layer MLP on the RNN hidden state trained with Adam at learning rate 10^−4^ against a Gaussian function of the true Euclidean distance to reward,

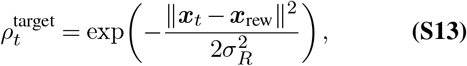

via

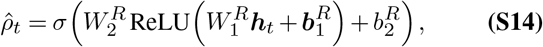

where 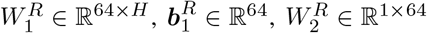 and 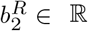 with *σ*_*R*_ = 0.15. Its mean-squared-error loss 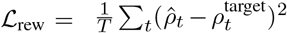 is added to the joint objective,

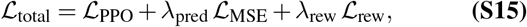

so that its gradient propagates through the decoder into the RNN weights. The target is computed directly from privileged agent position and the requested continuous reward coordinate at every step, it is dense supervision and not reward obtained from the environment, a temporal-difference error or an emergent dopamine ramp. We call it a reward-proximity teaching objective because the decoder error trains the recurrent representation to make proximity decodable. For this experiment we used the off-center reference coordinate ***x***_rew_ = (0.3, 0.3), with the training-grid approximation described above, set the RNN RL learning rate to 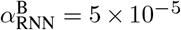, and used *λ*_pred_ = 0.3 and *λ*_rew_ = 1.5 for one Phase B cycle of 1000 episodes.

### Matched proximity-objective comparison and plasticity control

We separated recurrent plasticity from the presence of the proximity objective using networks that were identical except for the indicated manipulation.

#### Matched pre-learning state

Constructing the proximity decoder draws from the random-number generator, left uncorrected, this changes Phase A data, mini-batch order and replay trajectory. We therefore re-seeded immediately before Phase A, after decoder construction. Matched networks with and without the proximity objective then shared a byte-identical pre-RL place map and stored trajectory so post-learning differences can be attributed to the added objective within the implemented comparison.

#### Proximity objective versus no proximity objective

.Matched triples (no proximity objective, proximity objective, and near-rigid control) share a seed index and a byte-identical Phase A state. A triple is retained when all three networks pass the entropy-collapse gate on the periods listed under Gate periods below (Eq. (S17)). The near-rigid arm uses 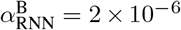, the plastic arms use 5 × 10^−5^, *λ*_pred_ = 0.3 and reward at (0.3, 0.3), differing only in whether the decoder and its objective are present (*λ*_rew_ = 1.5). Seed indices are not renumbered, so pairing by index remains valid. Of 109 launched triples, 55 are retained. Each run was analyzed with fixed-trajectory replay.

#### Stability guard

The replay probe is a 50,000-step continuous rollout, far longer than the 20-step training episodes, so a marginally unstable network can diverge during the probe even when its training curves look normal, producing activations orders of magnitude above the pre-RL scale and rendering distance-based measures meaningless (field peaks land on transient bursts). Rather than a magnitude threshold on neural activity, exclusion is handled by the fixed entropy-collapse gate defined below (Eq. (S17)), applied identically to every condition. It is the sole run-level map-quality exclusion criterion. Individual analyses additionally require enough eligible cells to define their statistic (at least 10 for paired shift estimates and 30 for reward-relative classification).

#### Plasticity control

The near-rigid arm of each matched triple 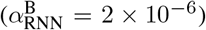 strongly restricts recurrent updates inside the RL block. Its reported replay snapshot is taken before the subsequent interleaved predictive block, whose learning rate remains 10^−4^. It therefore provides a low-plasticity control for the measured post-RL effect rather than a fully frozen network. We do not report a full dose– response: an earlier one-seed sweep was non-monotonic and confounded by map-integrity changes. The claim is restricted to the matched two-point contrast.

### Reward relocation

To test relocation, we used reward coordinates A = (0.3, 0.3) and B = (0.7, 0.7) for 1000 episodes each. Relocation simultaneously updated environmental reward, the controller’s supplied goal-displacement cue and the decoder’s proximity target, subject to this grid/continuous-coordinate distinction. A single pre-RL random-walk sequence was evaluated through baseline (*F* ^0^), post-A (*F* ^*A*^) and post-B (*F* ^*B*^) parameter snapshots. The standard cycle order was retained: *F* ^*A*^ was recorded immediately after the A RL block, followed by the A-policy interleaved predictive block, the reward was then changed and the B RL block produced *F* ^*B*^, before the B interleaved block. Consequently, *F* ^*A*^ → *F* ^*B*^ includes the intervening A-policy predictive update as well as learning in the B RL block, it is not an isolated B-only update. More generally, the analysis measures how stored parameters change when all goal-defining inputs are reassigned together, it does not test whether the network infers the new goal from reward experience alone. Tracking was quantified relative to the requested continuous coordinates A and B, using mean field-center distance and paired per-cell shifts *F* ^0^ → *F* ^*A*^ and *F* ^*A*^ → *F* ^*B*^.

### Spatial analysis and shift metrics

#### Rate maps and spatial information

Place-field analysis binned activity into a 32 × 32 grid and divided the summed activity in each bin by its occupancy, assigning zero to unvisited bins. Skaggs information was computed on this unsmoothed occupancy-normalized map (45). Field peaks and centers of mass were computed on a separate copy smoothed with a Gaussian kernel (*σ* = 1.5 bins) and rescaled to [0, 1]. Spatial selectivity used the Skaggs information score,

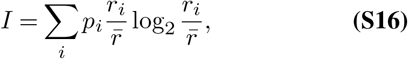

where *p*_*i*_ is the occupancy probability in bin *i, r*_*i*_ its mean activity, and 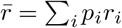 the occupancy-weighted mean activity. This single definition is used by every model analysis. The recurrent activations are continuous rather than spikes, so “bits/spike” in thresholds and figure labels is the conventional name for this normalized information index, not a literal spike-count unit.

#### Cell inclusion

Cells enter a paired contrast only if they exceed a Skaggs threshold of 0.3 bits/spike in *both* periods of that contrast. The threshold is 0.15 for relocation and reward omission. Each relocation step gates cells in the two snapshots being compared, the omission analysis follows cells through three snapshots and therefore requires the threshold in all three. Panels that display a single example network apply a looser 0.05 bits/spike gate, so that the displayed field distributions are not restricted to the most selective cells Every per-network population panel uses the gates above. Fig. S1f and Fig. S2b report sensitivity to this threshold. Example networks are selected objectively as the retained networks closest to the cohort median over all reported per-network shifts (peak and center of mass).

#### Behavioral and replay contrasts

The behavioral contrast compares the random-walk trajectory recorded before reward learning with the goal-directed probe recorded after it, so parameters and trajectory both differ, as in recordings made before and after learning. Fixed-trajectory replay instead compares the same random-walk trajectory evaluated through the two parameter sets. The goal-directed probe consists of short episodes: on each goal arrival the agent is moved to a random start and the recurrent state is reset. The five steps following each reset or segment start, which primarily reflect the reset transient, were excluded from goal-directed rate maps and from the occupancy-matched estimator.

#### Field location and sign convention

Field centers were taken as the center of mass of the smoothed, normalized rate map. Because a border-aligned or diffuse field can have its center of mass near the arena center, we additionally report the field *peak*, defined as the argmax of the smoothed map. With an off-center reward the peak metric is not pulled toward the arena center by this geometric symmetrization, and we report both measures throughout. Shift is

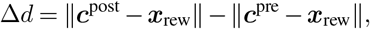

so Δ*d <* 0 means movement toward reward and Δ*d >* 0 means movement away.

#### Statistical unit and uncertainty

For population-level model claims, the independent unit is the trained network: cell-level shifts are first averaged within each network, and tests are then performed across network means. Paired tests compare networks with the same seed or cells from the same network, as specified. Pooled-cell distributions are descriptive unless a caption explicitly identifies a within-network cell-level test. Statistical significance used the two-sided Wilcoxon signed-rank test except where a one-sided, directionally predicted test is stated. Across-network values are reported as mean, standard deviation (s.d.) and standard error of the mean 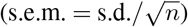; figure error bars show s.e.m. unless stated otherwise. Unit-level tuning classifications use Benjamini–Hochberg correction as described below, the distinct network-level contrasts address separate pre-specified questions and are reported without a family-wise correction. As a robustness check against diffuse or weakly tuned units, we recomputed shifts while progressively raising the Skaggs threshold, the toward-reward shift is essentially flat under this gating (Fig. S2b).

### Entropy-collapse seed gate

A minority of networks converge to a degenerate solution in which all field peaks collapse into a small region of the arena making field-center displacement meaningless. We exclude these with a fixed gate applied identically to every condition. For each period entering an analysis we compute the Shannon entropy (in bits) of the distribution of field peaks over the 32 × 32 grid, restricted to cells with Skaggs information at least 0.3, and require

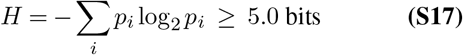

in *every* period used by that analysis. A network failing in any period is dropped from that analysis entirely. This is the only run-level exclusion based on map quality. The entropy gate uses the 0.3 cell threshold for all families. After network selection, relocation and omission effect estimates use their analysis-specific 0.15 paired/triple cell gates described above. Analysis-specific minimum cell counts are stated with their corresponding methods. The gate is computed with the same rate maps as the analyses (occupancy-weighted spatial information; goal-directed periods without the steps that follow each reset).

#### Gate periods

For matched triples, each plastic arm must pass on the post-RL random-walk and goal-directed probes and on both fixed-replay passes, the near-rigid arm must pass on both fixed-replay passes, and a seed is retained only if all three arms pass. Relocation networks must pass on the three replay snapshots and the three goal-directed periods used in the relocation analyses. Omission networks must pass on the baseline, learned and post-omission replay snapshots. These family-level gates keep the cohort fixed across the related comparisons rather than allowing each panel to select a different subset.

Networks were trained until at least 50 per family passed an earlier version of the gate. Seed indices are never renumbered and failed runs are retained, so launched-versus-passed counts remain recoverable. The threshold of 5.0 bits is the highest single value that retains at least 50 networks in every family, giving 55 of 109 launched paired triples, 52 of 72 launched relocation networks and 51 of 67 launched omission networks. We verified that the choice of threshold does not drive any result by recomputing every reported effect and contrast for thresholds from 3.5 to 6.5 bits (Fig. S2a). Between 3.5 and 5.5 bits (63 to 50 paired triples, 55 to 50 relocation networks and 55 to 46 omission networks), every effect and contrast kept its sign and remained significant (*p <* 0.05). At 6 and 6.5 bits, where 39 and 22 paired triples remain, every effect kept its sign and remained significant except the secondary comparison of behavioral shifts with and without proximity teaching (*p* = 0.11 and 0.46).

### Reward-distance and goal-referenced bearing analysis

For each timestep we computed distance to the active reward, the allocentric bearing *ψ*_*t*_ = atan2(*y*_rew_ − *y*_*t*_, *x*_rew_ − *x*_*t*_), and the egocentric reward bearing *φ*^rew^ = atan2(sin(*ψ*_*t*_ − *θ*_*t*_), cos(*ψ*_*t*_ − *θ*_*t*_)), where *θ*_*t*_ is heading. The analyzed activity was that of the 512 recurrent hidden units, controller latents were not used. Timesteps in a one-bin margin inside the outermost reachable position along each wall were excluded to reduce boundary-induced directional structure.

Distance modulation was the Pearson correlation between activity and negative reward distance, so positive values indicate greater activity near reward. To reject ordinary place fields that correlate with distance only because they occupy one angular sector, we also divided the data into six iso-distance rings and eight allocentric angular sectors and required approximate angular uniformity on fully sampled rings (at least 20 samples per sector). Egocentric bearing used 16 angular bins, each unit’s activity-weighted circular tuning curve gave a preferred bearing *φ*^∗,rew^ and mean resultant length (MRL). An allocentric head-direction control used the eight exact heading values.

For distance, goal-bearing and head-direction scores, the null distribution came from 500 independent cyclic shifts of the relevant behavioral covariate relative to activity. Empirical *p*-values were corrected across the 512 units with the Benjamini–Hochberg procedure at *q* ≤ 0.05 (46). Classification additionally required an absolute distance correlation of at least 0.1 for distance cells and an MRL of at least 0.1 for goal-direction and head-direction cells; distance cells also required a within-ring angular MRL below 0.15. For relocation we compared *φ*^∗,rew^ between phases in units passing the criterion in both and contrasted the circular-mean change with Δ_geom_, the rotation expected if tuning remained allocentric when the reference reward changed from A to B. Because the active reward enters this analysis as a covariate, the test establishes goal-referenced alignment but not the source of goal information.

### Occupancy-matched estimator

To test whether the fixed-trajectory dissociation could be approximated from behavioral recordings, we built an estimator using position, heading and activity from a pre-learning random walk and a post-learning goal-directed trajectory. Positions were binned on the same 32 × 32 grid, heading takes eight exact values. We defined joint (position bin, heading) strata, removed five steps after each hidden-state reset and subsampled both periods to the smaller count in each stratum, repeating this *K* = 20 times. We compared unmatched, position-matched, joint position × heading-matched and inverse-propensity-weighted variants. Skaggs information and paired-cell inclusion were recomputed from each matched map. Networks remained in the predeclared 55-triple cohort selected from the unmodified maps, no second entropy filter was applied after stochastic subsampling. Validation used fixed-trajectory replay as the reference across the 55 paired networks with and without the proximity objective under two post-period variants. The specificity control used a same-parameter contrast whose reference shift is zero. The ordering criterion asked whether matched behavioral estimates recover the paired ordering between objective conditions seen under fixed-trajectory replay.

### Reward-omission and proximity-withdrawal protocol

Starting from a network trained on reward A with the proximity objective, we continued training for a matched 1000-episode cycle in which environmental reward was set to zero: no goal bonus, progress term or step penalty. Goal-triggered episode termination and the controller’s goal-displacement cue remained active. Simultaneously, the proximity loss was removed (*λ*_rew_ → 0), so it no longer updated either the decoder or RNN, the frozen decoder was still evaluated to report how proximity information in its recurrent input changed. The manipulation therefore withdraws environmental reinforcement and proximity teaching while preserving two other sources of goal information, it does not simulate inference of reward-expectation extinction. Phase A, run seeds and rewarded-cycle hyperparameters matched the proximity condition, subject to the omission simulator’s recurrent-state indexing exception described above. As in relocation, the learned snapshot was recorded immediately after the rewarded RL block, the standard rewarded-policy interleaved predictive block then ran before the omission block and the withdrawn snapshot was recorded before the next interleaved block. The learned-to-withdrawn contrast therefore includes that intervening rewarded-policy predictive update as well as the omission RL block. The same stored trajectory was evaluated through baseline, learned and post-omission parameter snapshots. Networks were retained when they passed the entropy gate (Eq. (S17)) in all three periods; 51 of the 67 launched networks are retained. Near-reward enrichment is the fraction of field peaks within 0.15 of reward A. Reliability is the Pearson correlation between rate maps built from alternating 1000-step blocks of the replayed trajectory over bins visited in both block sets, near- and far-reward cells are split at 0.30 from A by their post-learning field peak.

### Reward-relative classification

To test for a subpopulation that preserves tuning relative to reward rather than the allocentric map, following Sosa et al. (35), we computed two per-cell correlations across relocation: *r*_place_ between allocentric maps and *r*_reward_ between reward-centered maps over their overlapping window. Because a stable allocentric field can have nonzero *r*_reward_, classification used the contrast between correlations and a cell-identity shuffle null. A unit was reward-relative if *r*_reward_ *> r*_place_ and *r*_reward_ exceeded the shuffle’s 95th percentile, place-anchored under the converse rule, and unclassified otherwise. Cells entered if their Skaggs information (occupancy-weighted) was at least 0.15 in both periods, networks with fewer than 30 such cells were excluded. Reward-centered maps covered ±0.3 around each reward, and each shuffle null used 2000 random cell-pair draws per network, discarding draws in which both members were the same cell. We report both goal-directed trajectories and fixed-trajectory evaluation, treating the latter as primary because sampling is identical. For the public recordings of Sosa et al. (35) we used their peak-distance criterion with a circular-roll null.

### Published-data reanalyses

The comparisons with recordings are post hoc tests of correspondence, not fits: no model parameter was estimated from either experimental dataset. For reward omission, we used the published source-data workbook of Krishnan et al. (34). We preserved the aggregation unit supplied for each measurement (animal for map reliability and field fractions, axon for VTA–CA1 fluorescence), normalized track position to [0, 1], and treated the end of the 200-cm track as the reward side. Start, middle and end regions span 0–50, 50–150 and 150–200 cm, respectively. Model and experimental quantities were plotted together only after each had been normalized as specified in the corresponding caption.

For reward relocation, we reanalyzed all seven within-environment switch sessions from two mice in DANDI 001361 (35); sessions that also changed environment were excluded. Calcium-event maps were restricted to running speeds of at least 2 cm/s, occupancy-normalized in 5-cm bins on the 450-cm circular track, and circularly smoothed with *σ* = 1 bin. We omitted the first 10 trials of each session and the first 5 trials after the reward switch. A cell entered if its spatial information exceeded the 95th percentile of 100 circular-shift surrogates before or after the switch and it was active in both periods; sessions required at least 30 such cells. A recorded cell was called reward-relative when its pre/post reward-centered peak discrepancy was less than 50 cm and below the fifth percentile of 500 circular rolls of its post-switch map. For the occupancy control, frames were matched within position-bin × speed-tercile strata over 10 repetitions; bins with less than 1 s occupancy per condition were treated as missing before circular interpolation. Session means are the plotted experimental units. Because the seven sessions come from only two mice, these comparisons are descriptive and do not establish population-level equivalence with the model.

**Table S1.** Hyperparameters for all experiments.

| Parameter | Value | Phase |
| --- | --- | --- |
| Hidden dimension $H$ | 512 | A, B |
| Observation dimension $M$ | 24 | A, B |
| Grid resolution | $32 \times 32$ | A, B |
| Trajectory length $T$ | 100 records (99 transitions) | A/interleave |
| Number of trajectories | 1000 | A |
| Training epochs (Phase A) | 100 | A |
| L1 regularization $\lambda_{L1}$ | $10^{-4}$ | A/interleave |
| Learning rate (Phase A) | $10^{-3}$ | A |
| Optimizer (Phase A) | RMSProp | A |
| Batch size | 64 | A |
| Activation bias init $\beta_0$ | -0.2 | A, B |
| RL episodes per cycle | 1000 | B |
| Number of cycles $N_{\text{cyc}}$ | 1 (replay), 2 (relocation) | B |
| Interleaved predictive epochs $E_{\text{int}}$ | 20 | B |
| Discount $\gamma$ | 0.99 | B |
| PPO clip $\epsilon$ | 0.1 | B |
| PPO epochs per episode | 4 | B |
| Random warm-up moves per episode | 10 | B |
| Entropy coefficient $c_2$ | 0.02 | B |
| Value loss coefficient $c_1$ | 0.5 | B |
| Advantage estimate | discounted return - value; boot-strap at truncation | B |
| Goal bonus / shaping / step penalty | +15 / 20 / -0.05 | B |
| Controller LR $\alpha_{\text{ctrl}}$ | $10^{-4}$ | B |
| Proximity-decoder LR | $10^{-4}$ | B |
| RNN RL LR $\alpha_{\text{RNN}}^{\text{B}}$, main | $5 \times 10^{-5}$ | B |
| RNN RL LR, rigid control | $2 \times 10^{-6}$ | B |
| Prediction weight $\lambda_{\text{pred}}$ | 0.3 (all conditions) | B |
| Online prediction regularizer | next-observation MSE (no L1) | B |
| Max episode length $T_{\text{max}}$ | 20 | B |
| Goal-probe logit divisor / segments | 0.3 / 50 | Analysis |
| Proximity-objective weight $\lambda_{\text{rew}}$ | 1.5 | B |
| Proximity-target width $\sigma_R$ | 0.15 | B |
| Reward reference $\mathbf{x}_{\text{rew}}$ | (0.3, 0.3) requested;<br>(0.3125, 0.3125) training grid | B |
| Skaggs threshold | 0.3 (0.15, relocation/omission)<br>bits/spike | Analysis |
| Entropy-collapse gate | 5.0 bits | Analysis |
| Rate map smoothing $\sigma$ | 1.5 bins | Analysis |
| Cycle goal-directed / final goal-directed / replay / post-cycle random-walk steps | 5000 / 50000 / 50000 / 10000 | Analysis |

### Hyperparameter summary

All hyperparameters are summarized in Table S1.

## Supplementary results

The Supplementary results are organized by the question each analysis resolves. S1–S3 show network-level distributions and robustness for the behavioral, replay and relocation claims. S4 asks whether reward-distance and goal-bearing tuning survives controls for sampling and reference frame. S5 and S6 expand the comparisons with reward-omission and reward-relocation recordings. S7 tests whether behavioral matching can substitute for fixed-trajectory replay, and S8 compares the recurrent maps with a trajectory-derived successor representation.

Unless stated otherwise, representational shifts use fixed-trajectory replay, field peaks and the paired place-cell gates defined in Methods; behavioral contrasts are labeled explicitly. A “paired” or “surviving” cell is one that passes the place-cell criterion in every period entering that contrast. An additional, unshown robustness pipeline using a stricter place-cell gate (Skaggs ≥ 1.0) and Gaussian-mixture field centers gave the same qualitative conclusions.

### S1. Map quality and behavioral shift, network by network

This section asks whether the behavioral concentration toward reward is broadly distributed across networks or driven by a small subset, and whether spatial selectivity survives Phase B. Because every headline population number is a mean over networks, we report the per-network values behind each one rather than relying on a representative example. Fig. S1 shows, for the 55 retained networks without proximity teaching, that the spatial map is preserved through Phase B and that the behavioral shift toward reward holds network by network, for field peaks and centers of mass and across place-cell thresholds.

### S2. Robustness of the main-text statistics and example-network distributions

This section separates robustness of the across-network conclusions from the cell-level distribution shown for the example network. Panel a varies the network-level entropy gate, panel b varies the cell-level Skaggs gate, and panel c shows the proximity-teaching example network from Fig. 4a,b (not the no-proximity example used in Figs. 2 and 3). Across Skaggs thresholds from 0.05 to 1 bits, the fixed-trajectory shift with proximity teaching stayed between −0.062 and −0.064 for field peaks, and the paired comparison with networks without proximity teaching remained significant (*p* ≤ 0.009 at every threshold; Fig. S2b), arguing against an artifact from weakly tuned units.

The network shown in Fig. 4a,b also exhibited the population-level effect at the level of individual cells (Fig. S2c). On the identical replayed trajectory, mean distance to reward decreased by 0.066 for field peaks and 0.036 for centers of mass, with shifts toward reward in 59% and 64% of eligible cells, respectively. These distributions illustrate the example network; the matched across-network plasticity test is shown in Fig. 4d.

### S3. Relocation, network by network

This section asks whether both stages of relocation, baseline to reward A, then reward A to reward B are consistent across networks and across definitions of field location. Fig. S3 gives the per-network statistics for the 52 retained networks, for field peaks and centers of mass. Both measures move toward A and then toward B, they differ in that field peaks move less toward B than toward A, whereas centers of mass move comparably toward both.

### S4. Reward-distance and goal-bearing analyses

This section asks whether apparent reward-vector tuning reflects a stable reference frame or the trajectories on which activity was sampled. Distance and bearing are both coupled to spatial sampling. We classified reward-distance cells only when activity both correlated with distance and remained approximately uniform around fully sampled iso-distance rings. Even with this control, distance-cell prevalence and goal-bearing tuning were much greater on goal-directed trajectories than on the replayed random walk (Fig. S4a). These measures therefore cannot establish a goal-centered code from tuning strength alone.

Relocation provides a more direct reference-frame test. A fully goal-relative code predicts no change in preferred goal bearing when the reward moves, whereas fixed allocentric tuning predicts a trajectory-dependent rotation. The observed change on goal-directed trajectories lay between these predictions (Fig. S4b); on the replayed trajectory, both the observed and allocentric-predicted changes were small, making that comparison inconclusive (Fig. S4c). The analyzed activity is the recurrent hidden state, not the controller latent, and receives no goal vector during fixed replay. These results therefore indicate partial goal-referenced tuning rather than a complete goal-vector code (33, 40).

### S5. Reward omission: reliability, proximity decoding and behavior

This section separates three consequences of withdrawing environmental reinforcement and proximity teaching: what the frozen decoder can read out, how stable the replayed maps remain, and how quickly reward-directed behavior stops. Fig. S5 expands the model–data comparison of Fig. 6. The proximity decoder, whose weights were fixed after learning, lost its spatial contrast because its recurrent input changed (Fig. S5a). In the model, split-half map reliability on the replayed trajectory fell from 0.692 (s.d. 0.127, s.e.m. 0.018) to 0.616 (s.d. 0.107, s.e.m. 0.015; *p* = 4.1 × 10^−4^) after withdrawal. The decline was confined to cells far from the former reward (0.732 → 0.615, *p* = 3.6 × 10^−6^); reliability near the reward had already fallen during learning and did not decline further (0.596 → 0.622, *p* = 0.21). In the recordings, by contrast, reliability declined in every track region, including the reward region (34) (Fig. S5b). In both model and mice, behavior directed at the former reward stopped within tens of episodes or a few laps (Fig. S5c,d).

### S6. CA1 reward-relocation reanalysis

This section shows which parts of the model–data comparison are stable to the reward-relative classification rule and to occupancy matching, and which are limited by temporal resolution. Fig. S6 shows the reanalyzed recordings of Sosa et al. (35) in more detail. Licking moves to the new reward zone within a few trials of the switch (Fig. S6a). Place-anchored cells keep their field position across the switch, whereas reward-relative cells move with the reward zone (Fig. S6b). The reward-relative fraction depends on the classification criterion: a looser rule based only on distance to reward classifies 22.3% (s.d. 2.5, s.e.m. 0.9) of recorded cells as reward-relative (Fig. S6c). Licking shifts to the new zone earlier than the estimated transfer of place-field density in most sessions, but the map estimate cannot resolve transfers faster than five trials (Fig. S6d,e). Occupancy matching leaves the field shift essentially unchanged on this track, where successive laps already follow nearly the same route (Fig. S6f). Because the seven sessions come from two animals, the session-level uncertainty shown here describes the sampled sessions rather than independent animal-level replication.

### S7. An occupancy-matched estimator: validation and its path-dependence limit

This section asks whether post hoc matching of behavioral samples can recover the parameter-dependent shift when a literal replay control is unavailable. Fig. S7 validates the behavioral estimator against fixed-trajectory replay. Position×heading matching brings the population mean closer to the reference (Fig. S7a) and removes a same-parameter artifact (Fig. S7b). Per-network agreement remains weak (*r* = −0.06 without the proximity objective and *r* = 0.21 with it; Fig. S7c). Moreover, the matched behavioral estimate is more negative without proximity teaching than with it (*p* = 4.6 × 10^−4^), opposite to the fixed-replay ordering (*p* = 1.8 × 10^−3^; Fig. S7d). Matching is therefore useful for reducing a known bias, but it is not an empirical substitute for holding the entire recurrent input history fixed.

### S8. Comparison with the successor representation

This section asks which parts of the behavioral field shift are predicted by a representation built directly from the observed transition statistics. We compared the recurrent fields with a successor representation (SR) estimated from the trajectories of each of the 55 matched networks (Fig. S8). The SR of a trajectory is *M* = (*I* − *γT*)^−1^, with *T* the empirical transition matrix between the 30 × 30 interior bins of the arena and *γ* = 0.99; the exploratory SR is that of the random walk, and the goal-directed SR is built from each network’s own goaldirected trajectory. Row *i* of *M* is the expected discounted future occupancy from state *i*; column *j* is the successor field of state *j*, the analogue of a place field (20, 21). Panel a uses centers of mass to compare field centroids with SR centroids, whereas panel b uses peaks, as in the primary replay analysis.

Under the goal-directed policy, expected future occupancy concentrated strongly toward reward (mean distance of occupancy centers to reward 0.34 → 0.14, s.e.m. 0.004, without proximity teaching; 0.34 → 0.15, s.e.m. 0.007, with it). Successor fields did not: their centers moved slightly away from reward (0.34 → 0.35, s.e.m. 0.003; *p <* 10^−3^ in both conditions), reflecting their backward skew along goal-directed paths, whereas the centers of the recurrent fields moved toward reward from exploration before learning to goal-directed behavior after it (0.33 → 0.21 and 0.33 → 0.23). For each place cell, the SR predicts the direction of its behavioral shift as the change in the successor-field center at the cell’s initial location. Observed and predicted directions were weakly aligned (61% and 58% of cells within 90°; pernetwork *V* = ⟨cos *θ*⟩ = 0.16, s.e.m. 0.03, *p* = 2.3 × 10^−6^, and 0.10, s.e.m. 0.03, *p* = 1.9 × 10^−4^), so changes in occupancy account for part of the direction of the behavioral shift. Under fixed-trajectory replay, by contrast, the trajectory-derived SR is unchanged by construction, whereas the recurrent fields moved toward the reward (−0.039, s.e.m. 0.007, and − 0.062, s.e.m. 0.004) more than toward control locations at the other corners of the inner square (between −0.022 and +0.016). This comparison is specific to the estimator implemented here and is not a claim that all successor-feature or reward-modulated SR models predict zero change.

**Fig. S1.**
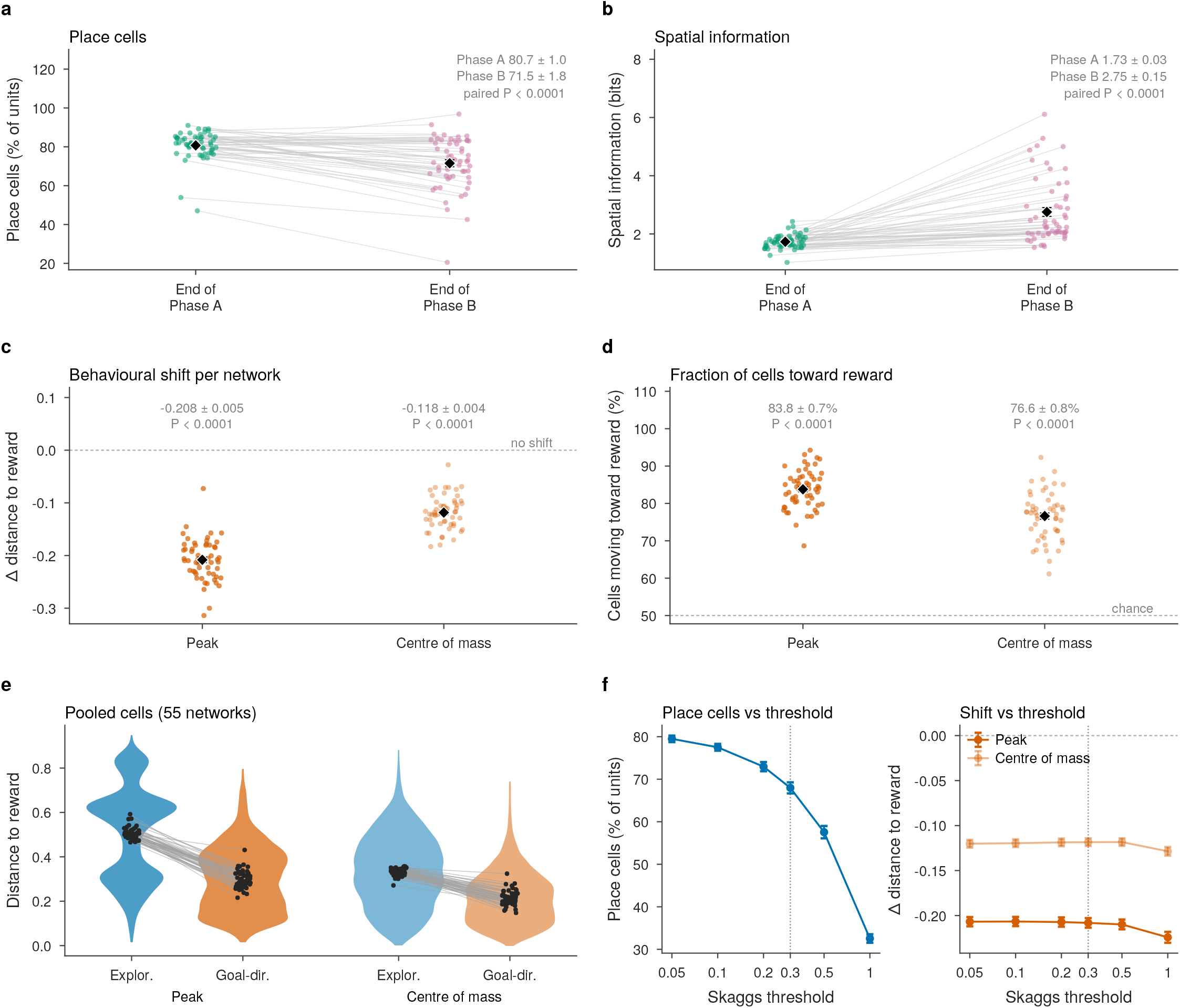
Map quality and behavioral shift, network by network. Across the 55 retained networks without proximity teaching used in the main text, the spatial map is preserved through reward learning (Phase B) and place fields concentrate near reward during goal-directed behavior in every network. The behavioral shift compares exploration before learning with goal-directed behavior after learning, so both parameters and trajectories differ. Place cells have Skaggs information ≥ 0.3 bits/spike in the periods compared. Each point is one network; black diamonds and error bars show the mean *±* s.e.m. a) Percentage of the 512 recurrent units that are place cells at the end of Phase A and at the end of Phase B, measured on the same replayed trajectory; gray lines join each network’s two values. b) Mean Skaggs spatial information of those place cells at the end of Phase A and of Phase B. In a,b, annotations give mean *±* s.e.m. and a paired Wilcoxon test. c) Behavioral shift: mean change in distance to reward (arena units) from exploration before learning to goal-directed behavior after learning, for field peaks and centers of mass (negative, toward reward; dashed line, no shift). d) Percentage of each network’s cells whose field moved toward reward in the same contrast (dashed line, 50% expected by chance). In c,d, annotations give mean *±* s.e.m. and a Wilcoxon test against zero or 50%. e) Distance to reward (arena units) of every place cell pooled over the 55 networks (violins) on the exploratory and goal-directed probes, for peaks and centers of mass; points are network means, joined between probes. f) Sensitivity to the Skaggs threshold (dotted line, 0.3 bits/spike): percentage of units that are place cells in both probes (left) and the behavioral shift for peaks and centers of mass (right); mean *±* s.e.m. across networks.

**Fig. S2.**
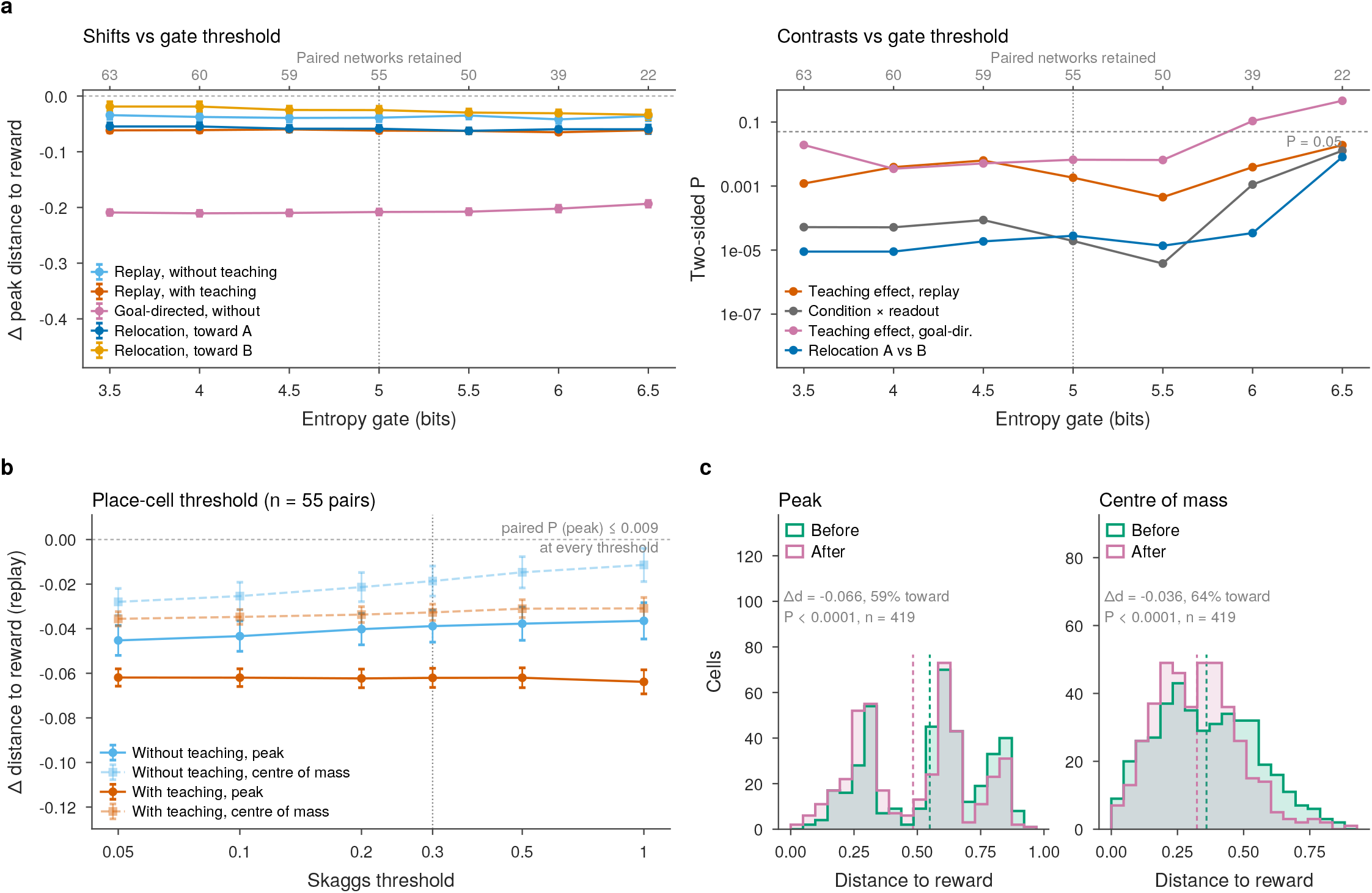
Robustness of the main-text statistics and example-network distributions. The main-text effects do not depend on the network-level entropy gate or the cell-level Skaggs threshold, and the proximity-teaching example network shows the reward-directed shift at the level of individual cells. a) Two plots show the dependence on the entropy-collapse gate, which excludes networks whose field peaks collapse into a small region of the arena (Eq. (S17)). Both share one x-axis (gate threshold); the dotted line marks the 5.0-bit threshold used throughout, and the numbers above each axis give the matched pairs retained at each threshold for the proximity-teaching comparisons. Left, effect sizes: field-peak shift (arena units; negative = toward reward; mean *±* s.e.m.) for five contrasts: replay without and with proximity teaching, goal-directed behavior without teaching, and relocation toward A and toward B. Right, significance: two-sided *P* of four main-text comparisons (dashed line, *P* = 0.05): the effect of proximity teaching on replay and on the goal-directed shift (paired by seed), the interaction between teaching and readout (replay versus goal-directed), and relocation shift toward A versus toward B. b) Dependence on the Skaggs place-cell threshold (dotted line, 0.3 bits/spike used throughout): fixed-trajectory shift without (blue) and with (orange) proximity teaching, for peaks (solid) and centers of mass (dashed), in the 55 matched pairs; mean *±* s.e.m. The paired comparison of peak shifts with and without teaching is significant at every threshold (*P* ≤ 0.009). c) Distance to reward (arena units) of the proximity-teaching example network’s place cells on the identical replayed trajectory before (green) and after (pink) learning, for field peaks and centers of mass (*n* = 419 cells with Skaggs information ≥ 0.05 bits/spike in both maps; dashed lines, means; annotations give the mean per-cell change, the percentage of cells moving toward reward and paired Wilcoxon tests).

**Fig. S3.**
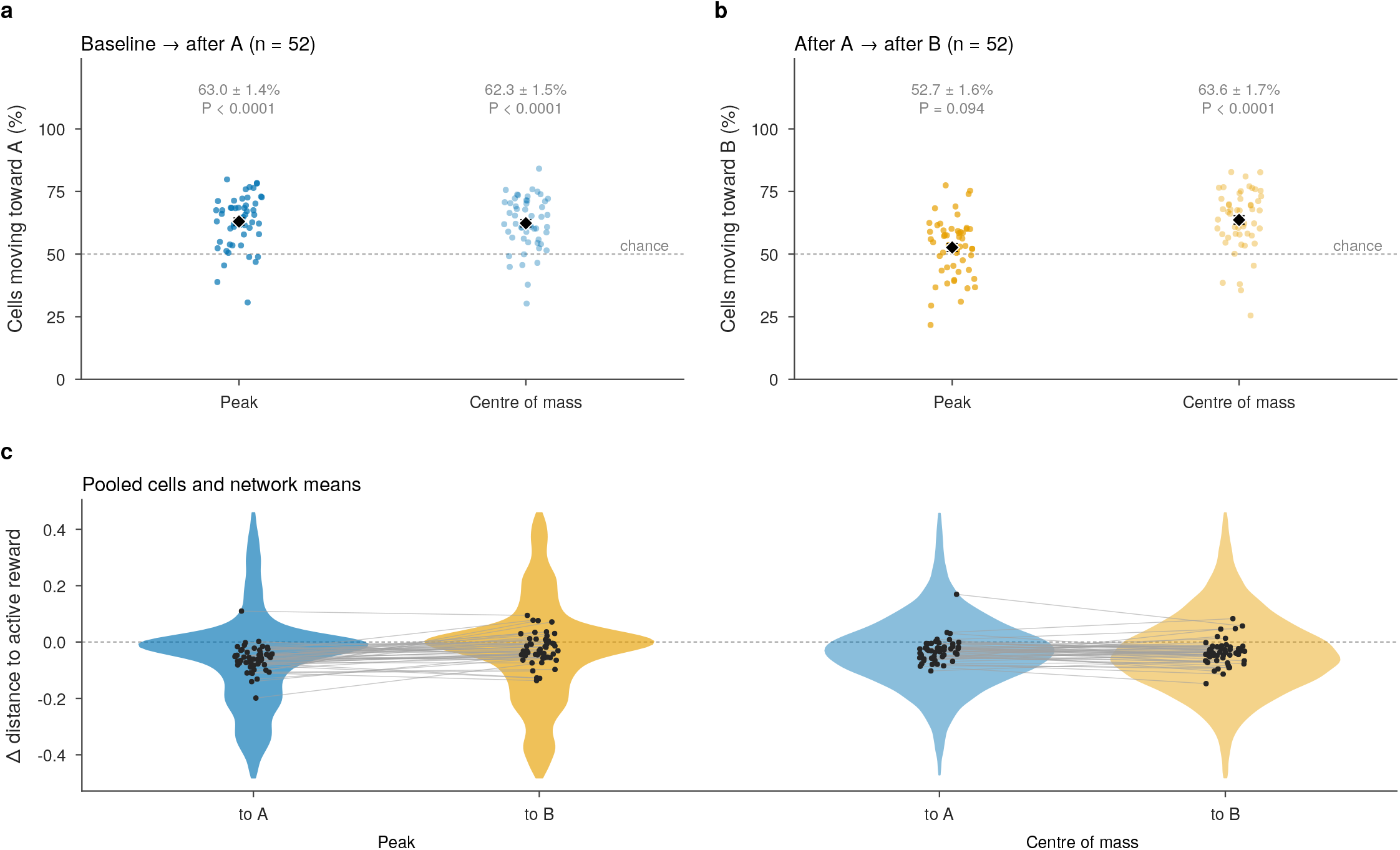
Relocation, network by network. Across the 52 retained relocation networks, mean field shifts are toward the active reward at both steps. The reward was learned at A = (0.3, 0.3) and then moved to B = (0.7, 0.7); fixed-trajectory replay passes the same stored exploratory observation–action trajectory through the baseline, after-A and after-B networks, and shifts were measured on cells that are place cells (Skaggs information ≥ 0.15 bits/spike) in both passes of each comparison. a) Percentage of each network’s place cells whose field moved toward A from baseline to after A, for field peaks and centers of mass (dashed line, 50% expected by chance); diamonds, mean *±* s.e.m.; annotations, mean *±* s.e.m. and a Wilcoxon test against 50%. b) The same for movement toward B from after A to after B; the fraction of cells moving toward B is above chance for centers of mass but not for peaks. c) Per-cell change in field distance to the active reward (arena units) for the two steps (negative, toward the reward), all cells pooled over networks (violins, central 98% of cells), with each network’s mean (points) joined between the two steps, for peaks and centers of mass.

**Fig. S4.**
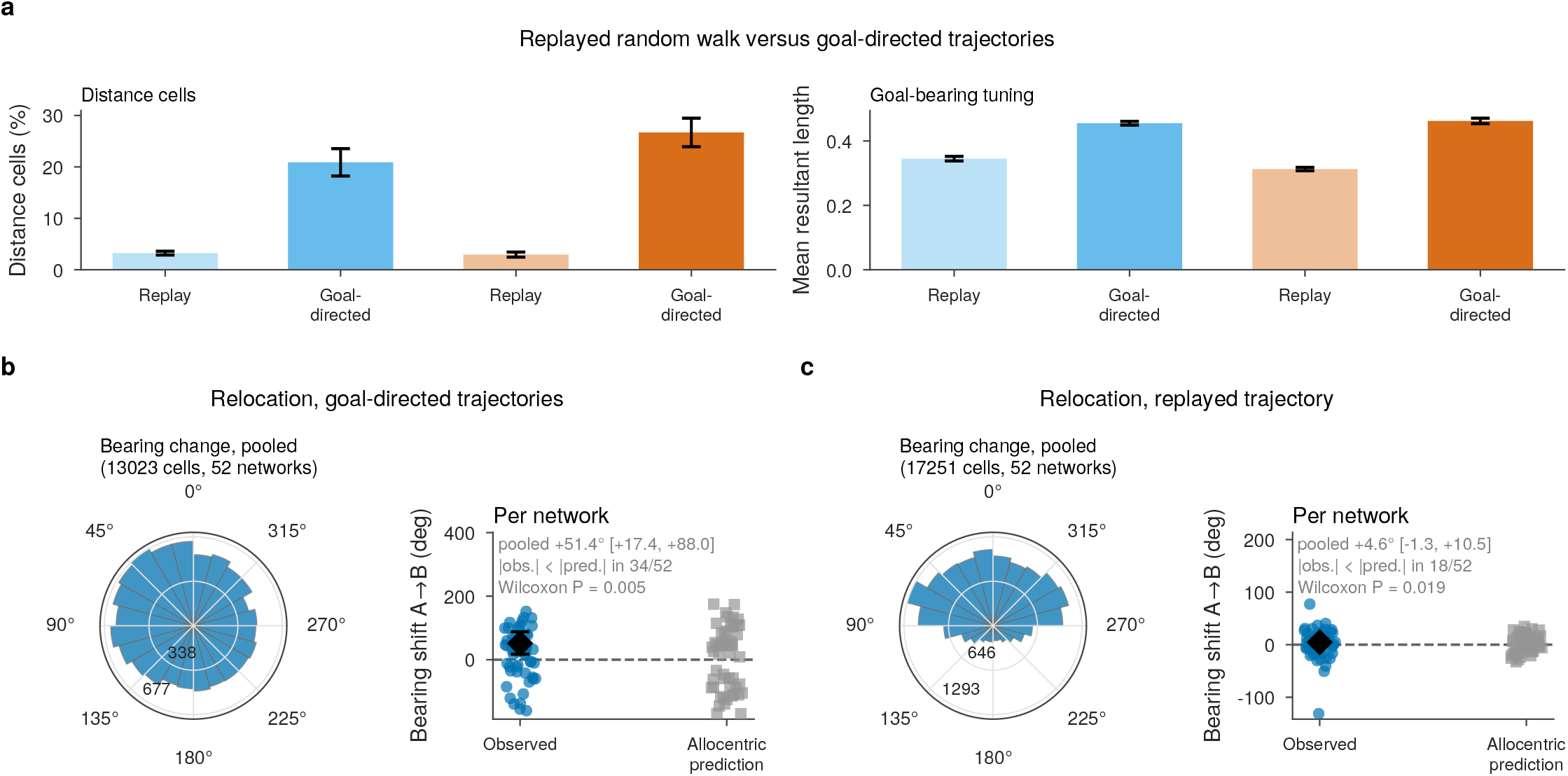
Reward-distance and goal-bearing measures depend on trajectory and reference frame. Reward-distance cells and goal-bearing tuning are stronger on goal-directed trajectories than under replay, and the preferred goal bearing changes after relocation on goal-directed trajectories but barely under replay. Analyses use the 512 recurrent units, excluding wall-adjacent timesteps; per-unit significance was assessed with cyclic-shift nulls, Benjamini–Hochberg correction and an effect-size floor. Fixed-trajectory replay passes one stored random-walk exploration trajectory through the network. a) Reward-distance cells (left, percentage of units) and mean goal-bearing tuning strength (right, mean resultant length) on fixed-trajectory replay and goal-directed trajectories, without (blue) and with (orange) proximity teaching; pale and dark bars denote replay and goal-directed sampling, respectively; error bars show mean *±* s.e.m. across 55 retained networks. b,c) Change in preferred goal bearing after relocation of the reward from A to B on goal-directed trajectories (b) and under fixed-trajectory replay (c) across 52 retained relocation networks. Left, rose plots pool all qualifying units (13,023 in b, 17,251 in c). Right, each network’s observed circular-mean change (blue) is compared with the rotation predicted by purely allocentric tuning (gray); diamonds show the pooled mean and network-resampled 95% confidence interval; annotations also give the number of networks whose observed shift is smaller in magnitude than the prediction (|obs.| *<* |pred.|) and a Wilcoxon test. The observed pooled changes were +51.4^°^ ([+17.4^°^, +88.0^°^]) during goal-directed behavior and +4.6^°^ ([−1.3^°^, +10.5^°^]) during replay.

**Fig. S5.**
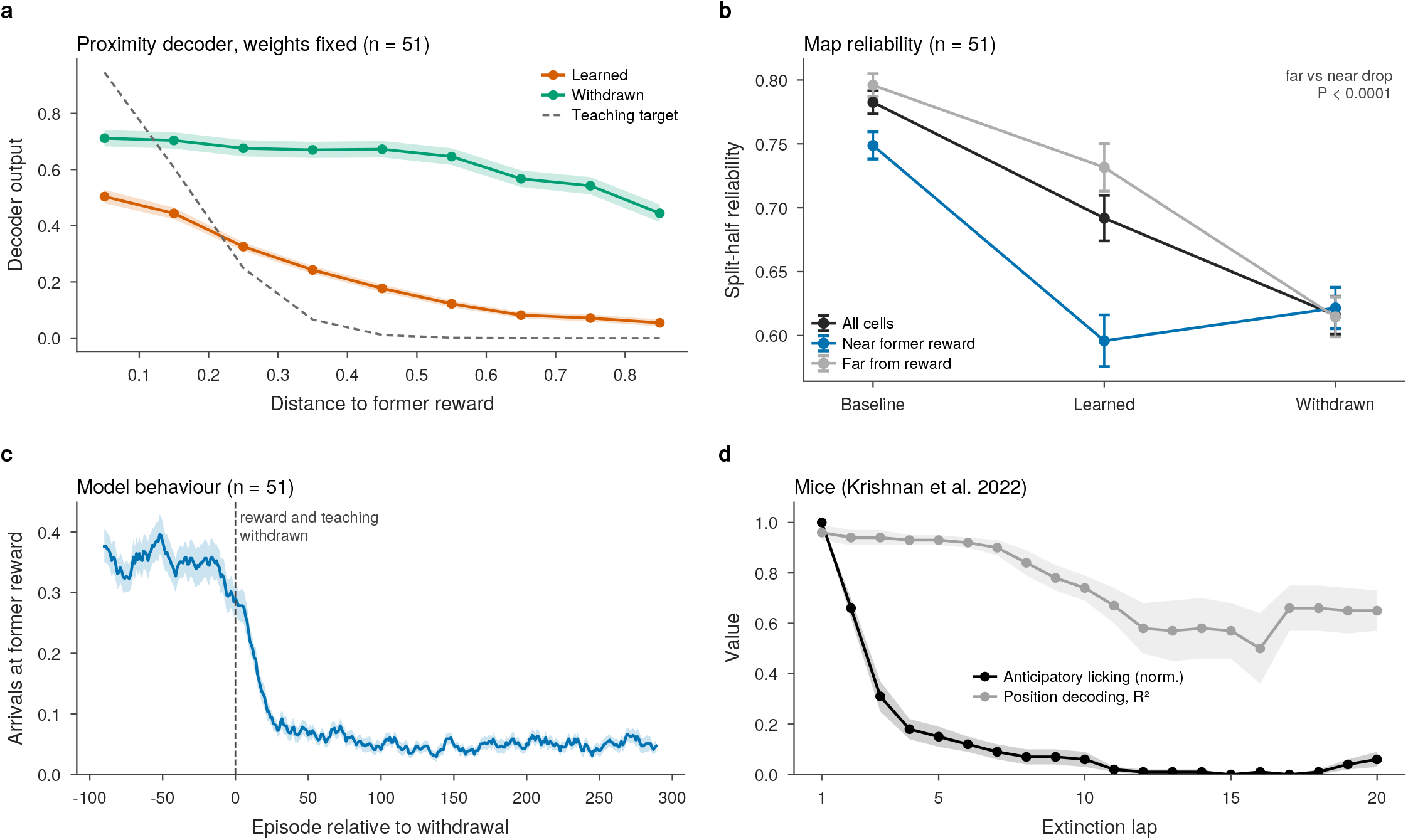
Reward omission: reliability, proximity decoding and behavior. Withdrawing reward and teaching flattens the model’s proximity readout, equalizes map reliability near and far from the former reward and reduces arrivals at the former reward, whereas in mice licking extinguishes faster than position decoding declines. Environmental reward and the proximity teaching signal are withdrawn together after learning with proximity teaching. a) Output of the proximity decoder, with weights fixed after learning, against distance to the former reward (arena units) on the fixed-trajectory replay, learned (orange) and withdrawn (green) (mean *±* s.e.m.), with its teaching target (dashed). b) Split-half map reliability on the fixed-trajectory replay through baseline, learned and withdrawn weights, for all place cells (black) and for cells near (≤ 0.15; blue) and far from (gray) the former reward (*n* = 51 networks; mean *±* s.e.m.). Near-reward reliability falls from baseline to learned, whereas far-from-reward reliability falls from learned to withdrawn; the *P* value is a paired test of the far versus near decline from learned to withdrawn. c) Model behavior: arrivals at the former reward per episode around withdrawal (dashed line) (10-episode running mean, mean *±* s.e.m. over networks). d) Mice across extinction laps: normalized anticipatory licking (black) and position-decoding accuracy (gray, *R*^2^) (mean *±* s.e.m., published source data of Krishnan et al. 2022, Fig. 1e).

**Fig. S6.**
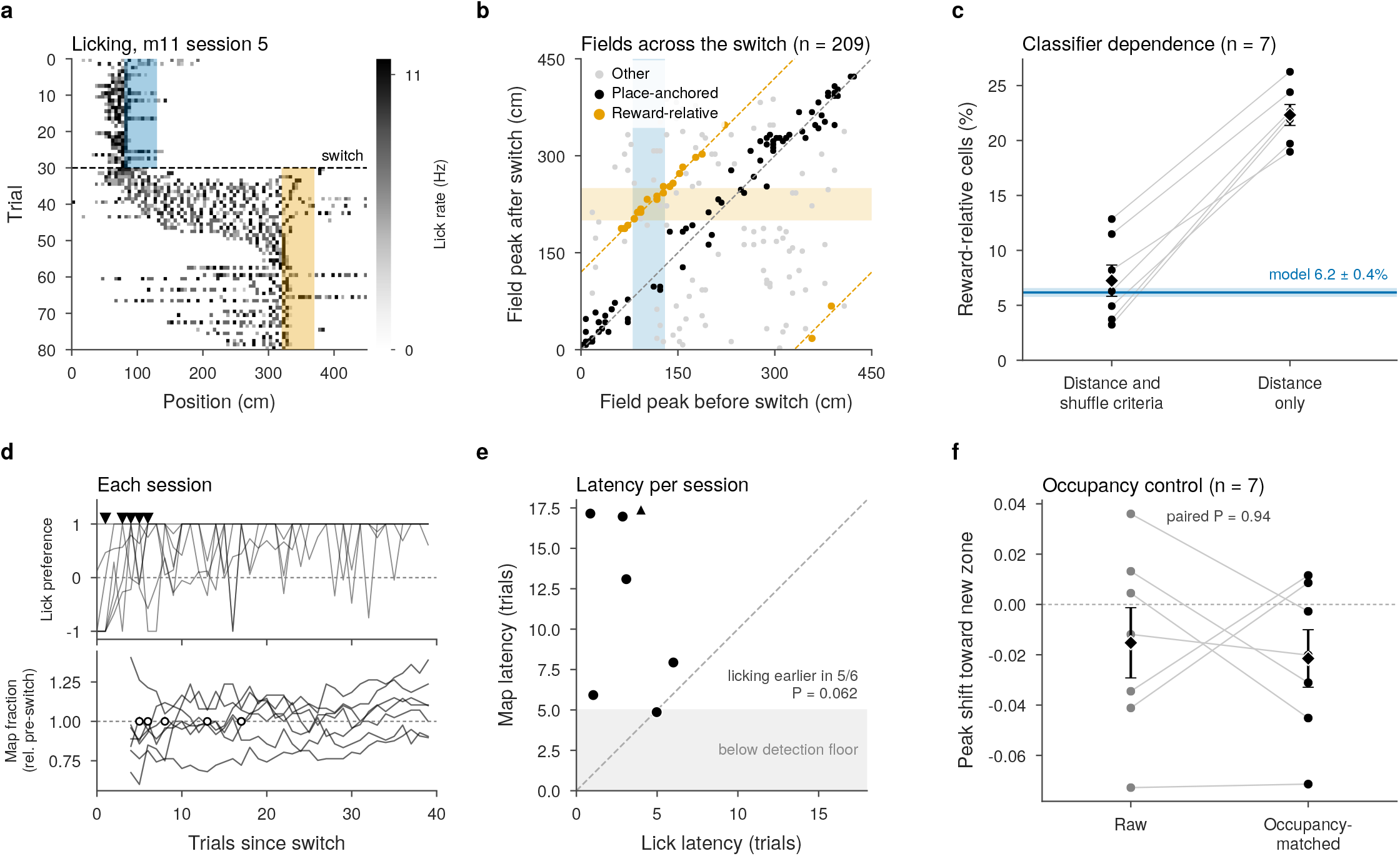
CA1 reward-relocation reanalysis. In the recordings, the reward-relative fraction is higher under the distance criterion alone than under the full criterion, and licking retargets earlier than the map in five of six decidable sessions. Recordings from Sosa et al. (35) (DANDI 001361; seven switch sessions, two mice). a) Lick rate by position and trial in one session (mouse m11, session 5; dashed line, switch); shading marks the reward zone before (blue) and after (orange) the switch. b) Field peak before against after the switch for the place cells of one session (*n* = 209 cells): place-anchored cells (black) lie on the identity line, reward-relative cells (orange) on the line offset by the zone shift (dashed lines); shading, reward zones. c) Reward-relative fraction per session under the full criterion (within 50 cm and beyond a shuffle control) and under the distance criterion alone, with the model mean *±* s.e.m. (blue band); diamonds, session mean *±* s.e.m. d) Each session’s lick preference for the new zone (top; triangles, lick latency) and map fraction relative to the pre-switch level (bottom; circles, map latency) after the switch; lines, sessions. e) Lick and map retargeting latency per session (points) (dashed line, equality; triangle, a session whose map never reached the pre-switch level). Licking is earlier in five of six decidable sessions (Wilcoxon *p* = 0.0625, the smallest attainable at this sample size); map latency cannot be detected below five trials (shaded). f) Raw and occupancy-matched shift of field peaks toward the new zone, per session (negative, toward); lines join each session’s two values; annotation, paired *P* value; diamonds, mean *±* s.e.m.

**Fig. S7.**
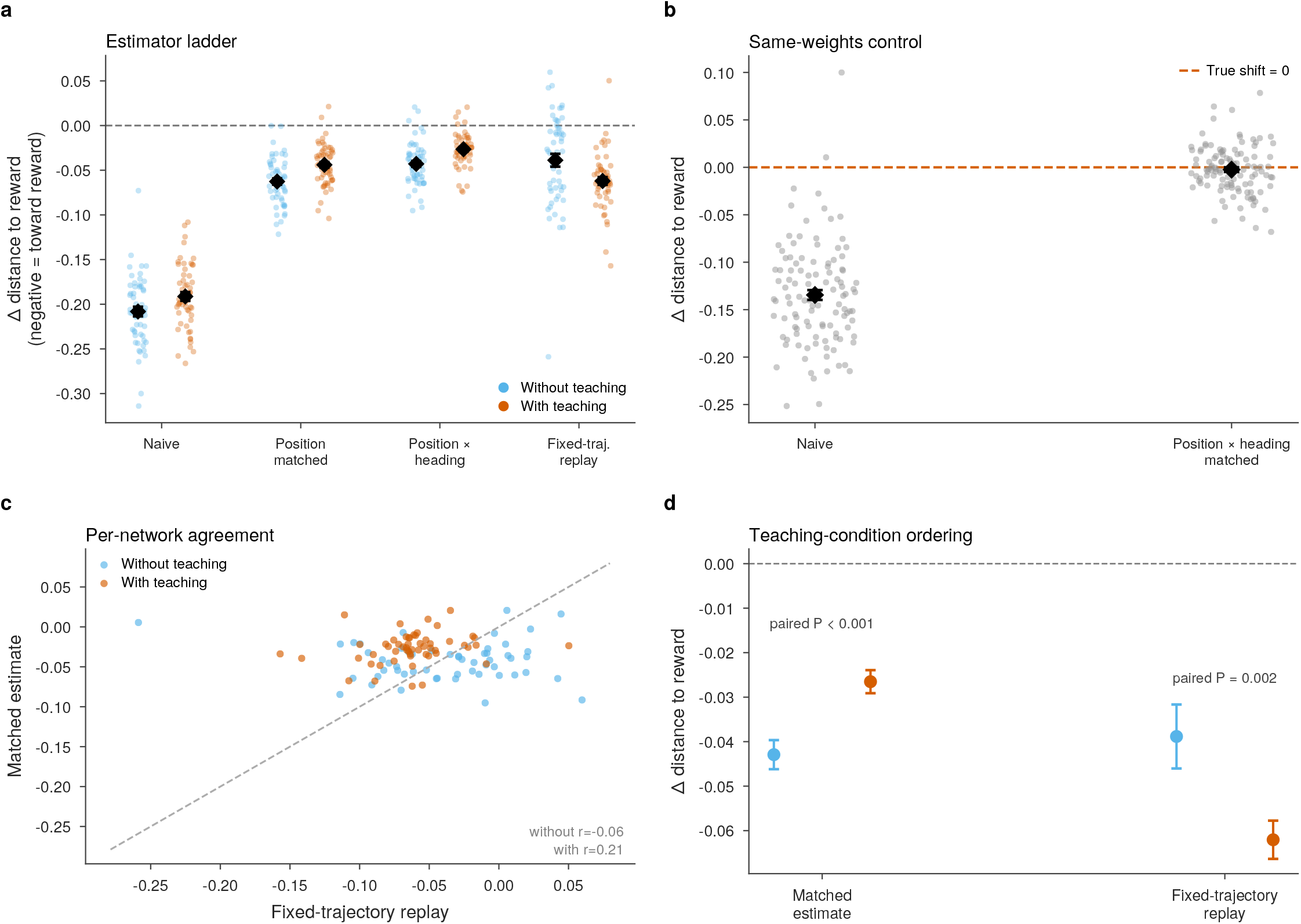
Occupancy matching versus fixed-trajectory replay. The matched estimator does not recover the ordering of the two teaching conditions that fixed-trajectory replay gives, and it agrees only weakly with replay network by network. The occupancy-matched estimator asks whether the fixed-trajectory shift can be recovered from behavioral data alone: activity from the pre-learning random walk and from the post-learning goal-directed trajectory is subsampled so that both periods contain the same number of timesteps in each position (and heading) bin before field shifts are measured. Each point is one network (*n* = 55 per condition); black diamonds and error bars show the mean *±* s.e.m.; blue, without and orange, with the proximity objective. a) Estimated change in field distance to reward (arena units; negative = toward reward), from left to right: the unmatched behavioral readout (naive), position-matched, position*×*heading-matched, and the fixed-trajectory replay that serves as ground truth. Matching removes most of the sampling bias of the naive readout. b) Specificity control: the random-walk and goal-directed probes run with the same post-learning weights, so the true shift is zero (dashed line). The naive readout reports a large spurious shift; the position*×*heading-matched estimator reports a shift close to zero. c) Per-network matched estimate against the fixed-trajectory shift, both conditions shown (blue, without; orange, with) (dashed line, identity); annotations give the correlation *r* for each condition; agreement for individual networks is weak. d) The pre-specified ordering test, paired by seed: mean shift with and without the proximity objective under the matched estimator and under fixed-trajectory replay. Only fixed-trajectory replay recovers the ordering of the two conditions (a larger shift with the objective); the matched estimator gives the reverse ordering; error bars, mean *±* s.e.m.; annotations, paired *P* values.

**Fig. S8.**
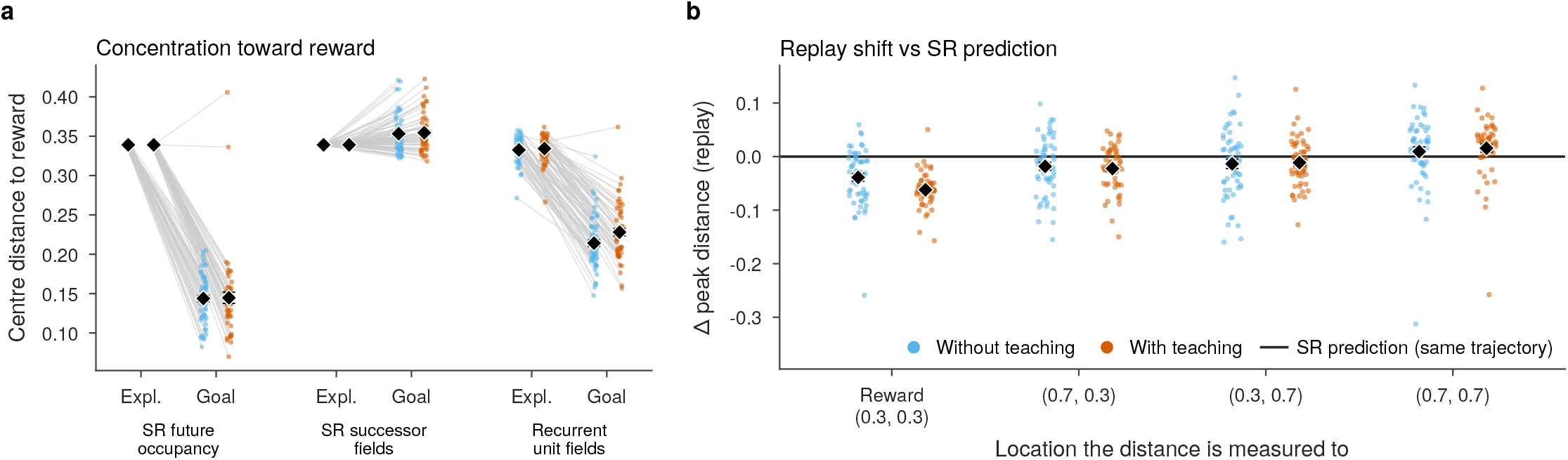
Comparison with the successor representation. Unlike the successor representation, the recurrent units shift toward the reward under fixed-trajectory replay. Both panels use the 55 matched networks without (blue) and with (orange) proximity teaching; each point is one network and diamonds show mean *±* s.e.m. a) Mean distance to reward (arena units) of the centers of SR expected future occupancy (rows of *M*), of SR successor fields (columns of *M*) and of the recurrent units’ place fields (center of mass, Skaggs ≥ 0.3 bits/spike), for the exploratory (Expl.) and goal-directed (Goal) trajectories; gray lines join a network’s two values. The exploratory SR is the same for every network. b) Fixed-trajectory replay shift of recurrent-field peaks, measured as the change in distance to the reward (0.3, 0.3) and to three control locations (0.7, 0.3), (0.3, 0.7) and (0.7, 0.7); the black line marks the SR prediction for an identical trajectory (zero).

